# Formation of a μ_₃_-oxo nucleophile enables efficient hydrolysis by a trinuclear metal center in Family II inorganic pyrophosphatase

**DOI:** 10.1101/2025.06.08.658471

**Authors:** Saki Maruoka, Yohei Kametani, Eisuke Magome, Hiroyuki Setoyama, Masahide Kawamoto, Masaki Horitani, Takamasa Teramoto, Yoshimitsu Kakuta, Yoshihito Shiota, Kazunari Yoshizawa, Keiichi Watanabe

**Author notes:** **Corresponding author:** Keiichi Watanabe.

## Abstract

Efficient catalysis by metalloproteins relies on precise spatial arrangement of metal ions and active-site residues. Family II inorganic pyrophosphatase (PPase) from *Shewanella* species features a trinuclear metal center and displays higher catalytic activity than binuclear counterparts. Here we elucidate its hydrolytic mechanism using X-ray crystal structure-based extended X-ray absorption fine structure (XCS-EXAFS), site-directed mutagenesis, and density functional theory (DFT) calculations. We identify a catalytic μ₃-oxo nucleophile, formed via proton transfer from a bridging μ₃-hydroxide to Asp14 and subsequent hydrogen-bond rearrangement to Asp72, as the key species in S_N_2-type hydrolysis. This conversion defines the rate-limiting step with an activation barrier of 15.5 kcal/mol. Molecular orbital analysis reveals that the trinuclear cluster promotes μ₃-oxo formation, aligns the nucleophile for attack, and stabilizes the transition state. The side-chain rotation of the conserved Asp14 is crucial for catalysis. Our results highlight how metalloenzymes exploit geometric and electronic tuning to achieve high reactivity through evolutionarily optimized architectures.

## Introduction

Nature employs highly sophisticated strategies to precisely position metal ions within enzyme active sites, thereby enabling accurate and efficient hydrolytic reactions. Many hydrolases utilize two divalent metal cations, such as Zn²⁺, Mg²⁺, Ni²⁺, Mn²⁺, or Fe²⁺, to coordinate a water-derived oxygen that functions as a nucleophile^1–3^. These structural adaptations underlie the remarkable specificity and catalytic efficiency observed in biological systems, supporting essential functions such as metabolism and regulation.

A prominent example is the cleavage of phosphoester (P–O) bonds involved in phosphate metabolism, particularly in pathways utilizing inorganic pyrophosphate (POP) and its derivatives^4,5^. Hydrolytic reactions of this type are catalyzed by phosphatases and pyrophosphatases, which are primarily classified by the nature of their nucleophiles and metal coordination modes^6^. Representative nucleophilic species include cysteine thiolates in tyrosine phosphatases (type A)^7,8^, serine alkoxides in PhoA-type alkaline phosphatases (type B)^9,10^, terminal hydroxides coordinated to a single metal ion as in purple acid phosphatases and 3’-5’ exonucleases (type C) ^11–13^, and bridging hydroxides stabilized by two metal ions as seen in phosphotriesterases and serine/threonine phosphatases (type D)^14–16^. A distinct μ₃-oxo species coordinated by three metal cations has been observed in PhoX-type alkaline phosphatases from Pseudomonas fluorescens (type E)^17^.

In this study, we focused on inorganic pyrophosphatases (PPases), a class of essential enzymes involved in intracellular phosphate metabolism (Fig. 1a)^18,19^. Previous X-ray crystallographic studies of Family II PPases from *Shewanella* sp. AS-11 (ShPPase) and *Bacillus subtilis* (BsPPase), complexed with a substrate analog imidodiphosphate (PNP), have revealed a unique trinuclear metal center. Each metal ion coordinates both a terminal oxygen of the electrophilic phosphate and a lone pair of the putative nucleophile, aligning them linearly with the scissile bond (Fig. 1b, c)^20,21^. However, it has remained unclear how the nucleophile— a μ₃-hydroxide with all three lone pairs engaged in metal coordination—can attack the electrophilic phosphorus center (Fig. 1b).

**Fig. 1.**
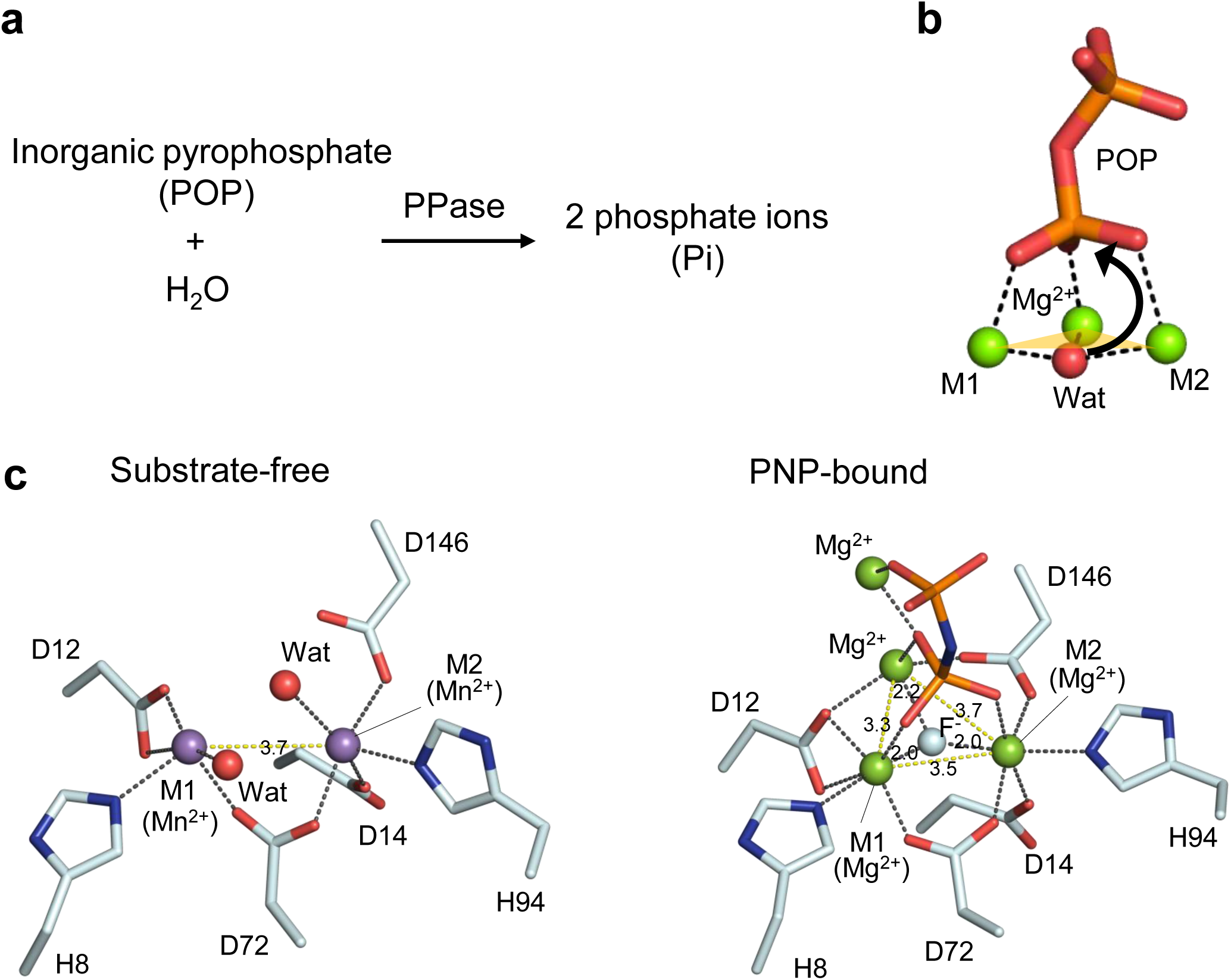
Catalytic function and active site structures of ShPPase. **a**, Schematic of the pyrophosphatase-catalysed hydrolysis reaction. **b**, Model depicting the proposed nucleophilic attack by a µ₃-OH^-^ species based on crystallographic data. Black dashed lines indicate coordination bonds; the yellow triangle denotes the plane defined by the trinuclear metal cluster. **c**, Coordination geometries of the metal ions in the active sites of the substrate-free Mn²⁺– ShPPase (PDB: 6LL7, left) and the PNP-complexed Mg²⁺–ShPPase (PDB: 6LL8, right). Amino acid residues and the substrate analogue PNP are shown as sticks; metal ions are represented as spheres. A five-coordinate geometry is observed at M1 and M2 in the substrate-free state, while a six-coordinate geometry is seen in the PNP-bound state. Mn, Mg, O, N, P, and H atoms are coloured purple, green, red, blue, orange, and white, respectively.

To address this question, we employed X-ray crystal structure-based extended X-ray absorption fine structure (XCS-EXAFS) spectroscopy, site-directed mutagenesis, and density functional theory (DFT) calculations using a trinuclear metal cluster model of the active site. Our mechanistic analysis reveals that, unlike the canonical μ-hydroxide-bridged binuclear mechanism (type D), hydrolysis by ShPPase proceeds via a distinct μ₃-oxo intermediate formed within the trinuclear metal center. Notably, the side chain of the conserved general base Asp14 undergoes a conformational rotation to accept a proton from the μ₃-hydroxide, forming the active μ₃-oxo nucleophile. This mechanistic feature provides a clear explanation for the higher catalytic efficiency of Family II PPases compared to their binuclear Family I counterparts^18,22^. Our findings advance the understanding of phosphoester bond cleavage by metalloenzymes and offer valuable insights for the design of biomimetic catalysts and the engineering of metalloenzyme functions.

## Results and discussion

### EXAFS analysis of substrate-free and PNP-bound ShPPase

In the substrate-free state, the active site of Family II ShPPase accommodates two divalent transition metal cations at the M1 and M2 sites^18,20^. Previous crystallographic studies have shown that Mn²⁺ ions occupy these positions in the substrate-free form (Mn²⁺-ShPPase, PDB: 6LL7) (Fig. 1c, left)^20^. In contrast, upon binding of the substrate analog PNP, an additional metal cation (Mg²⁺) is incorporated into the dinuclear center to generate a trinuclear metal site (Mg²⁺-ShPPase, PDB: 6LL8) (Fig. 1c, right)^20^. In this structure, the inhibitor F⁻ is positioned at the center of the metal cluster, and is presumed to be replaced by a nucleophilic OH⁻ species during catalysis. To elucidate how a μ₃-hydroxide species, lacking a free lone pair, could participate in nucleophilic attack on the phosphorus atom, we performed high-resolution EXAFS analysis (with 0.01 Å-level resolution) to determine the metal coordination geometry in solution ^23–26^.

The metal ions at the M1 and M2 sites of ShPPase are exchangeable in vitro, and previous studies have shown that hydrolytic activity is retained with various divalent transition metals^27^. For EXAFS measurements, we selected Zn²⁺-activated ShPPase (Zn²⁺-ShPPase), which exhibits both strong X-ray absorption (Fig. S1) and sufficient enzymatic activity (Table 1). Comparative EXAFS analyses of the substrate-free and PNP-bound forms revealed distinct differences in both XANES (X-ray absorption near-edge structure) and EXAFS spectra (Fig. 2). Notably, PNP binding led to electronic perturbation in the Zn²⁺ 3d orbitals, as reflected by an increased white-line intensity (Fig. 2a). Fourier-transform (FT) analysis showed a prominent first-shell peak at R + Δ = 1.65 Å, along with weaker second- and third-shell peaks at R + Δ = 2.5–4.0 Å, suggesting structural reorganization upon substrate binding (Fig. 2b).

**Fig. 2.**
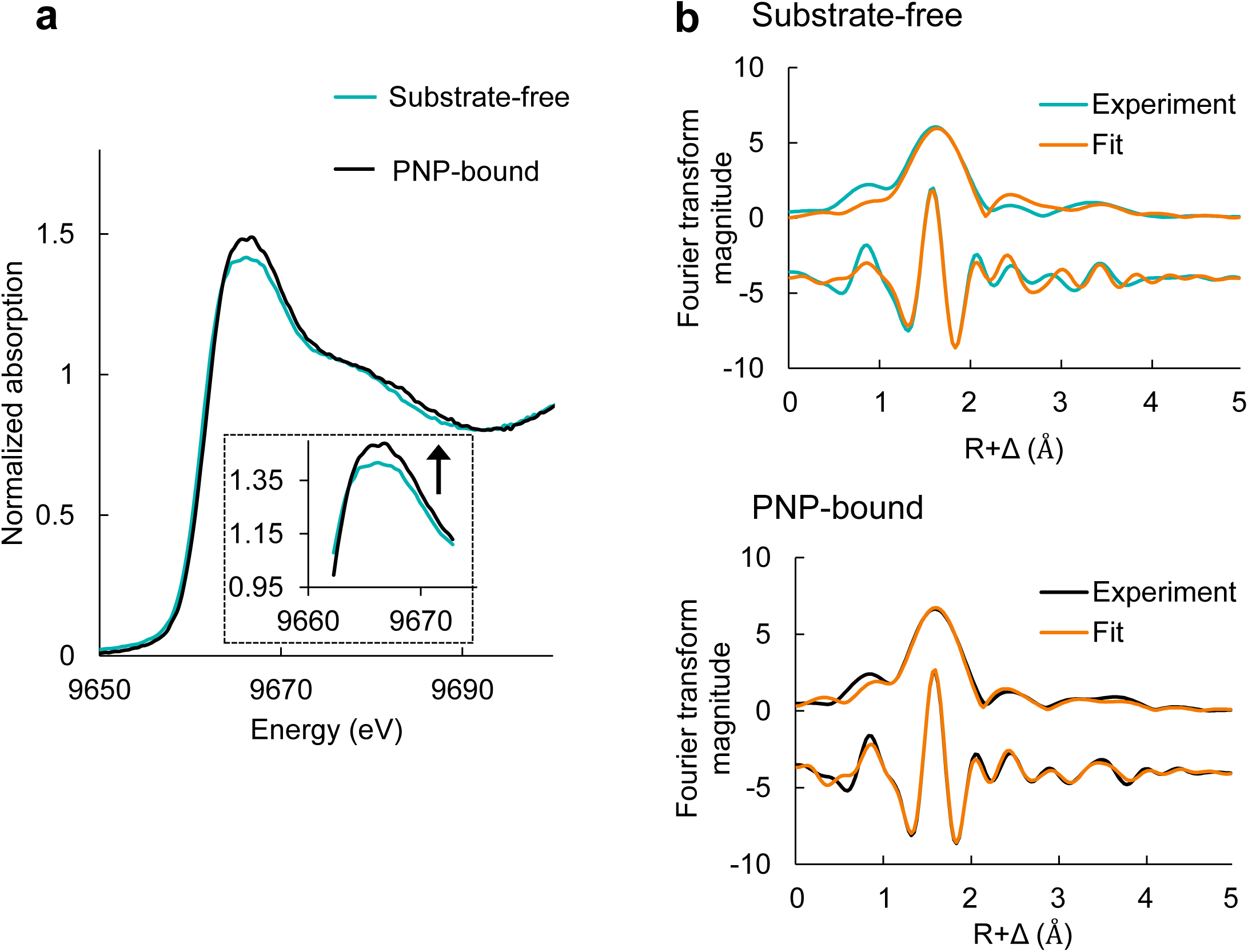
XAFS analysis of Zn²⁺–ShPPase. **a**, Zn K-edge XANES spectra of the substrate-free (light blue) and PNP-bound (black) forms. **b**, Fourier-transformed EXAFS spectra. Light blue and black lines correspond to experimental data for the substrate-free and PNP-bound forms, respectively; orange lines represent best-fit simulations.

**Table 1.** Kinetic parameters of Zn^2+^-ShPPases.

| Zn<sup>2+</sup>-ShPPase | *k<sub>cat</sub>* (s<sup>-1</sup>) | *K<sub>m</sub>* (μM) | *k<sub>cat</sub>*/*K<sub>m</sub>* (s<sup>-1</sup>μM<sup>-1</sup>) |
| --- | --- | --- | --- |
| Wild type | 99,800 ± 5,500 | 79 ± 18 | 1,260 |
| D14A | 19 ± 1.28 | 179 ± 37 | 0.11 |
| D14N | 34 ± 1.82 | 100 ± 20 | 0.34 |

Kinetic parameters determined from activity assays. Data are obtained with three times measurement.

Based on the Mn²⁺-ShPPase crystal structure, the first coordination shell likely corresponds to 4–5 oxygen and nitrogen atoms located approximately 2.5 Å from Zn²⁺ (Fig. 1c, 2b). Outer-shell features arise from single and multiple scattering paths involving histidine and aspartate residues, as well as the phosphorus atom of PNP, consistent with R values in the 3–5 Å range. The relatively high symmetry of Zn–Zn distances observed in the crystal structure allowed identification of four major single-scattering paths: Zn–O, Zn–N, Zn–C, and Zn–Zn (Fig. S2).

Curve fitting in k-space using parameters derived from the crystal structure revealed dynamic changes in the coordination environment upon PNP binding. Prior studies have demonstrated that metal substitution in the active site does not significantly perturb the overall structure^20,21,28^, validating our experimental approach. We performed the fitting analysis using constraint parameters as described in the Methods section^23^ (Table 2). The results showed that Zn²⁺ ions are predominantly five-coordinate in the substrate-free form, but transition to six-coordinate upon PNP binding. This change aligns with the crystallographic data and accurately reflects the altered coordination environment (Table 2, Fig. 1c). These findings highlight the sensitivity of EXAFS in capturing subtle changes in metal coordination geometry and support the use of this technique for probing metalloenzyme active sites under physiologically relevant conditions.

**Table 2.** Best fits for EXAFS data of Zn^2+^-ShPPases.

| ShPPase | Site | Absorber-Scatterer | N | R (Å) | $\sigma^2$ | E<sub>0</sub> | R<sub>factor</sub> |
| --- | --- | --- | --- | --- | --- | --- | --- |
| Substrate-free | M1 | Zn-O | 3.5 | 2.06 | 0.0075 | 9.999 | 0.784 |
|  |  | Zn-N | 1 | 2.56 |  |  |  |
|  |  | Zn-Zn | 1 | 3.97 |  |  |  |
|  | M2 | Zn-O | 3.5 | 2.05 | 0.0069 | 9.998 | 0.871 |
|  |  | Zn-N | 1 | 2.57 |  |  |  |
|  |  | Zn-Zn | 1 | 3.94 |  |  |  |
| PNP-bound | M1 | Zn-O | 4 | 2.02 | 0.0034 | 3.478 | 0.035 |
| | | Zn-O ($\mu_3$) | 1 | 1.82 | | | |
|  |  | Zn-N | 1 | 2.14 |  |  |  |
|  |  | Zn-P | 1 | 3.31 |  |  |  |
|  |  | Zn-Zn | 1 | 3.33 |  |  |  |
|  | M2 | Zn-O | 4 | 2.11 | 0.0126 | 5.684 | 0.029 |
| | | Zn-O ($\mu_3$) | 1 | 1.82 | | | |
|  |  | Zn-N | 1 | 2.13 |  |  |  |
|  |  | Zn-P | 1 | 3.27 |  |  |  |
|  |  | Zn-Zn | 1 | 3.32 |  |  |  |

Each row corresponds to a best fit for data obtained from measurements on an independently prepared set of samples. N, coordination number; R, interatomic distance (Å); $\sigma^2$ , meansquare deviation in R (the Debye-Waller factor) (Å<sup>2</sup>); E<sub>0</sub>, threshold energy shift (eV).

### Structural evidence for a μ₃-oxo nucleophile in the PNP-bound complex

EXAFS fitting revealed that the Zn–O (water) bond distance is significantly shortened from 2.06 Å in the substrate-free state to 1.82 Å in the PNP-bound complex, indicating a stronger coordination induced by substrate binding (Table 2). The Zn–Zn distance also decreased from 3.78 Å to 3.33 Å, suggesting substantial structural reorganization within the trinuclear metal center. Although subtle, these changes were clearly resolved by EXAFS, demonstrating its sensitivity to fundamental alterations in metal coordination environments at the active site.

A particularly noteworthy feature is the presence of a Zn–O(μ₃) bond at 1.82 Å, identified using an independent fitting parameter among the five oxygen atoms coordinating each Zn²⁺ ion. This result supports the assignment of a μ₃-oxo (O²⁻) species^24,25^ bridging two Zn²⁺ ions within the active site. The validity of this interpretation is further supported by comparison with the Fe–O distance of 1.80-1.83 Å observed for the bridging oxo group at a binuclear iron center of oxyhemerythrin^26^. The formation of a μ₃-oxo species implies deprotonation of a hydroxide ion to generate an oxo species, which plays a critical role in catalysis by facilitating nucleophilic attack on the phosphorus atom of inorganic pyrophosphate (POP). This structural transformation is therefore central to the enzymatic mechanism of phosphate bond cleavage.

### Construction and optimization of a computational model of the ShPPase active site

To further validate the structural insights obtained from EXAFS, we performed DFT-based mechanistic modeling using the high-resolution crystal structure of the PNP-bound complex (PDB ID: 6LL8). Prior to mechanistic analysis, we constructed and evaluated a computational model that included PNP, four metal ions, and a fluoride ion occupying the position corresponding to the nucleophilic water molecule. This model was optimized using DFT, yielding an average Zn–O bond distance of 2.13 Å, which is in good agreement with the 2.0 Å observed in the crystal structure, thus supporting the validity of the model. Based on this foundation, we constructed both a PNP-bound model (representative of the EXAFS experimental condition) and a model for the catalytic reaction with inorganic pyrophosphate (POP), by substituting specific atoms or ligands as detailed in the Methods section and Figs. S3–S6.

To better reproduce the enzyme environment used for EXAFS measurements, the crystal structure model was modified by replacing fluorine with hydroxide or oxygen and Mg²⁺ at the M1 and M2 sites with Zn²⁺. In these optimized models, the calculated Zn–O (μ₃) bond distances were 2.22 Å for the μ₃-hydroxide state and 1.94 Å for the μ₃-oxo state (Fig. S4). The shorter distance in the μ₃-oxo configuration supports the presence of an oxo ion (O²⁻), consistent with previous DFT studies of other metalloprotein cores^6,17^. Additionally, the Zn–Zn distances decreased from 3.68 Å in the μ₃-hydroxide state to 3.17 Å in the μ₃-oxo state, further supporting a compaction of the metal cluster upon formation of the oxo species. These computational results align closely with our EXAFS data and strongly support the conclusion that the μ₃-oxo species constitutes a critical structural element during catalysis (Fig. S7).

### Functional significance of Asp14 in μ₃-hydroxide deprotonation in ShPPase

The mechanism by which the μ₃-hydroxide species is deprotonated in ShPPase has remained unresolved. Although recent structural studies have accumulated detailed insights into Family II PPases, the process by which deprotonation occurs within the trinuclear metal center has not been fully elucidated. Conventional models, based on Family I PPases, have proposed that nucleophilic attack proceeds via a μ-hydroxide possessing a free lone pair^18,29^. However, in the highly stable trinuclear metal configuration of Family II PPases, formation of such a lone pair would require partial dissociation of at least one metal ion, introducing a substantial energetic barrier.

To explore potential general base candidates responsible for deprotonating the μ₃-hydroxide, we examined the X-ray crystal structure of the PNP-bound form (PDB: 6LL8). Four aspartate residues were found in close proximity to the μ₃-hydroxide moiety (Fig. 1c, right). Among them, three residues (Asp12, Asp72, and Asp146) coordinate metal ions via both side-chain carboxylate oxygens, contributing to the formation and stabilization of the trinuclear center. In contrast, Asp14 engages only one of its carboxylate oxygens (Oδ1) in metal coordination, leaving the other (Oδ2) available for potential involvement in proton abstraction. To test the functional role of Asp14, site-directed mutagenesis was performed. Both D14A and D14N variants exhibited drastic reductions in catalytic activity, with *k*_cat_/*K*_m_ values decreasing to approximately 1/4,000–1/10,000 of that of the wild-type enzyme (Table 1). In contrast, *K*_m_ values increased only modestly (approximately twofold), indicating that substrate binding remained largely intact (Table 1, Fig. S8). DFT calculations using a cluster model confirmed that the overall trinuclear architecture of the active site was maintained in these mutants (Fig. S9), ruling out structural disruption as the cause of activity loss. Furthermore, sequence homology analysis revealed that Asp14 is strictly conserved across all Family II PPases (Fig. S10), strongly supporting its essential role in μ₃-hydroxide deprotonation and nucleophile activation.

### S_N_2-type hydrolysis catalyzed by a μ₃-oxo species in the trinuclear metal active site

To elucidate the catalytic mechanism of inorganic pyrophosphate (POP) hydrolysis in Family II PPase, we performed DFT calculations based on structural models incorporating POP as the substrate (Figs. S5, S6). The reaction proceeds through seven discrete states, including three transition states (TS) and two intermediates (IM), capturing the key structural and energetic transformations leading to product formation (Fig. 3). The mechanism unfolds in three sequential stages:

**Fig. 3.**
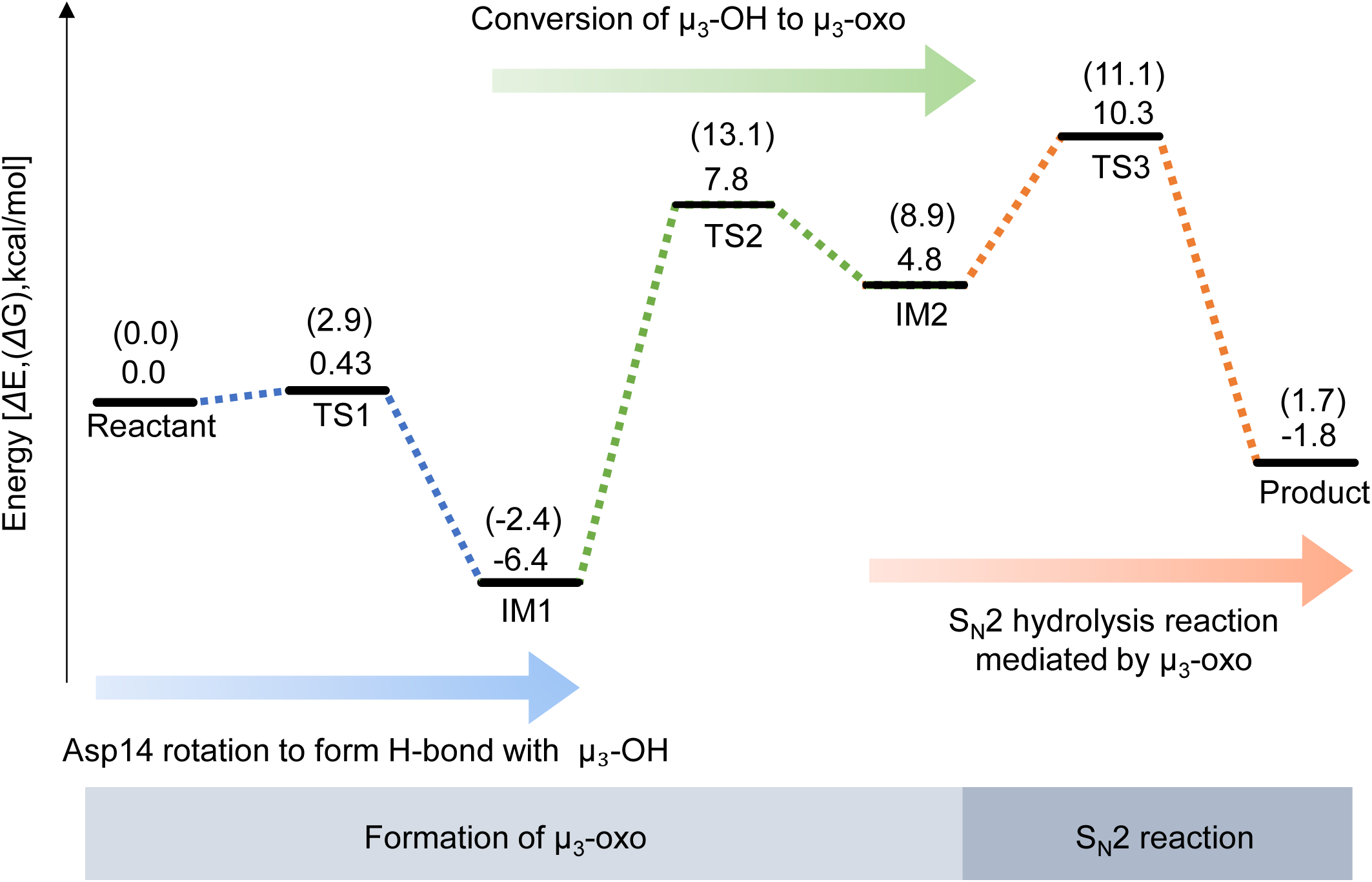
Free energy profile for the POP hydrolysis reaction in Zn²⁺–ShPPase. Gibbs free energy changes (in kcal mol⁻¹) are shown in brackets. The reaction pathway consists of Asp14 rotation to form H-bond with µ₃-OH (blue dashed line), Conversion of μ_3_-OH to μ_3_-oxo (green dashed line), and S_N_2 hydrolysis (orange dashed line).

Stage 1: Hydrogen bond formation via rotation of Asp14.

In the crystal structures (PDB: 6LL7, 6LL8), Asp14 forms hydrogen bonds with Ser116 and Asn117. To investigate how Asp14 mediates deprotonation of the μ₃-hydroxide species, we constructed a rotation model of Asp14 (Fig. S5). In the optimized reactant state, the Oδ1 atom of Asp14 coordinates a Zn²⁺ ion, while Oδ2 forms hydrogen bonds with the Oγ and N atoms of Ser116 at distances of 2.06 Å and 2.10 Å, respectively (Fig. 4). At this stage, the Oδ2–H(μ₃-hydroxide) distance is 3.70 Å.

**Fig. 4.**
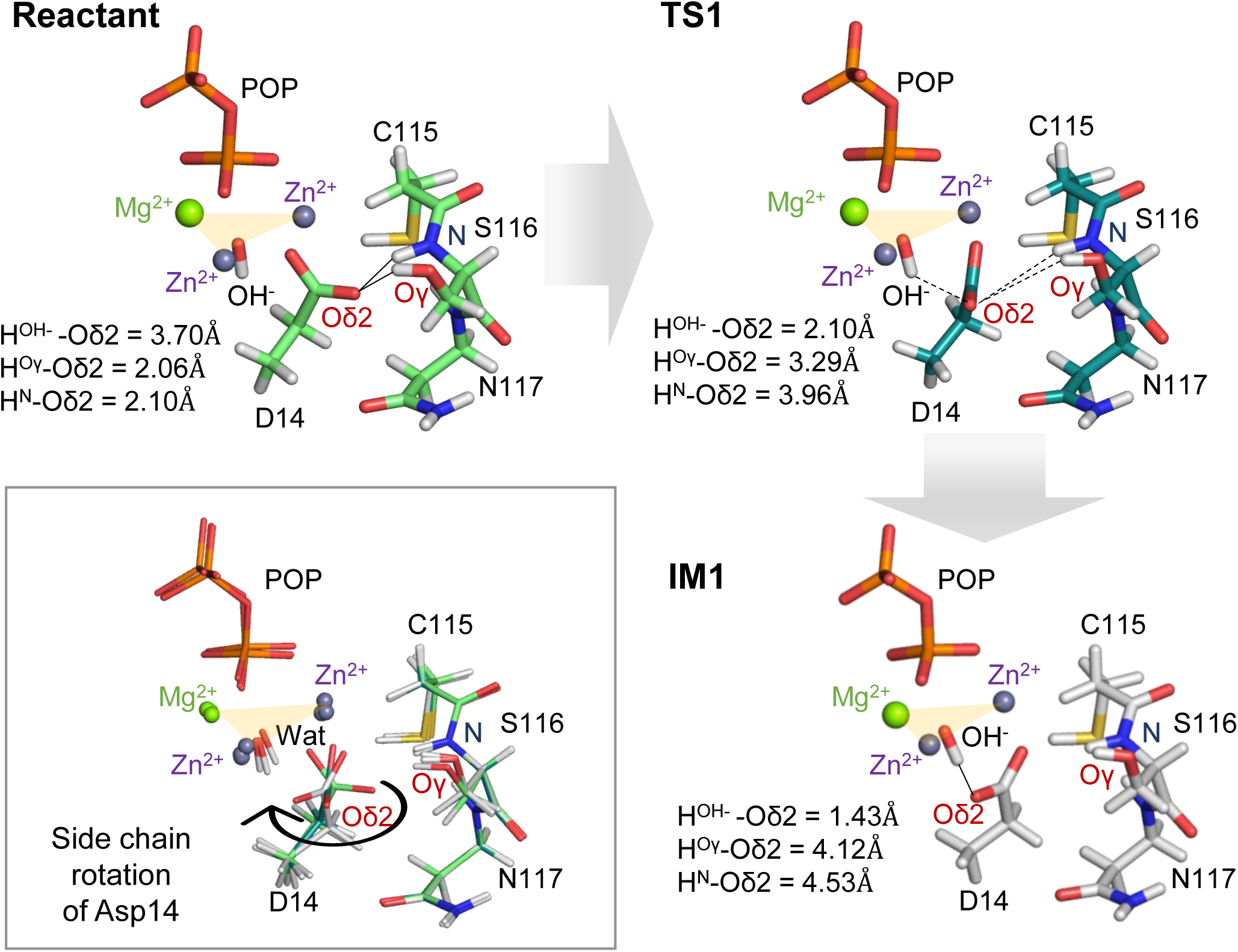
Side chain rotation of Asp14 in Zn²⁺–ShPPase. Optimized structures of the reactant state (light green, upper left), the transition state TS1 (dark green, upper right), and the intermediate IM1 (white, lower right) are shown as sticks. Superpositions of Cα-aligned structures are boxed in grey. Only the active-site region is displayed to emphasize key residue rearrangements. For a complete model of Asp14 rotation, see Fig. S5. Zn, Mg, O, N, P, and H atoms are coloured slate purple, green, red, blue, orange, and white, respectively.

Upon rotation of the Asp14 side chain, a hydrogen bond can form between Oδ2 and the μ₃-hydroxide, leading from the reactant state to the IM1 intermediate via TS1 (Fig. 4, S5). Crystal structures indicate sufficient spatial allowance around Asp14 to accommodate this rotation. The transition from the reactant to TS1 requires only a small activation barrier of 2.9 kcal/mol (ΔG), stabilizing the IM1 structure (Figs. 3). During this process, the original hydrogen bonds are disrupted, and a new hydrogen bond forms between Oδ2 and H(μ₃-hydroxide) at 1.43 Å (Fig. 4), priming the system for nucleophile activation.

Stage 2: Conversion of μ₃-hydroxide to μ₃-oxo.

This stage requires further rotation of the Asp14 side chain (Figs. 5, S6). We defined a “rotation angle” of Asp14 (0° at IM1; positive away from the μ₃-oxo toward Asp72 Oδ2 and negative toward the μ₃-oxo) to quantify this movement.

**Fig. 5.**
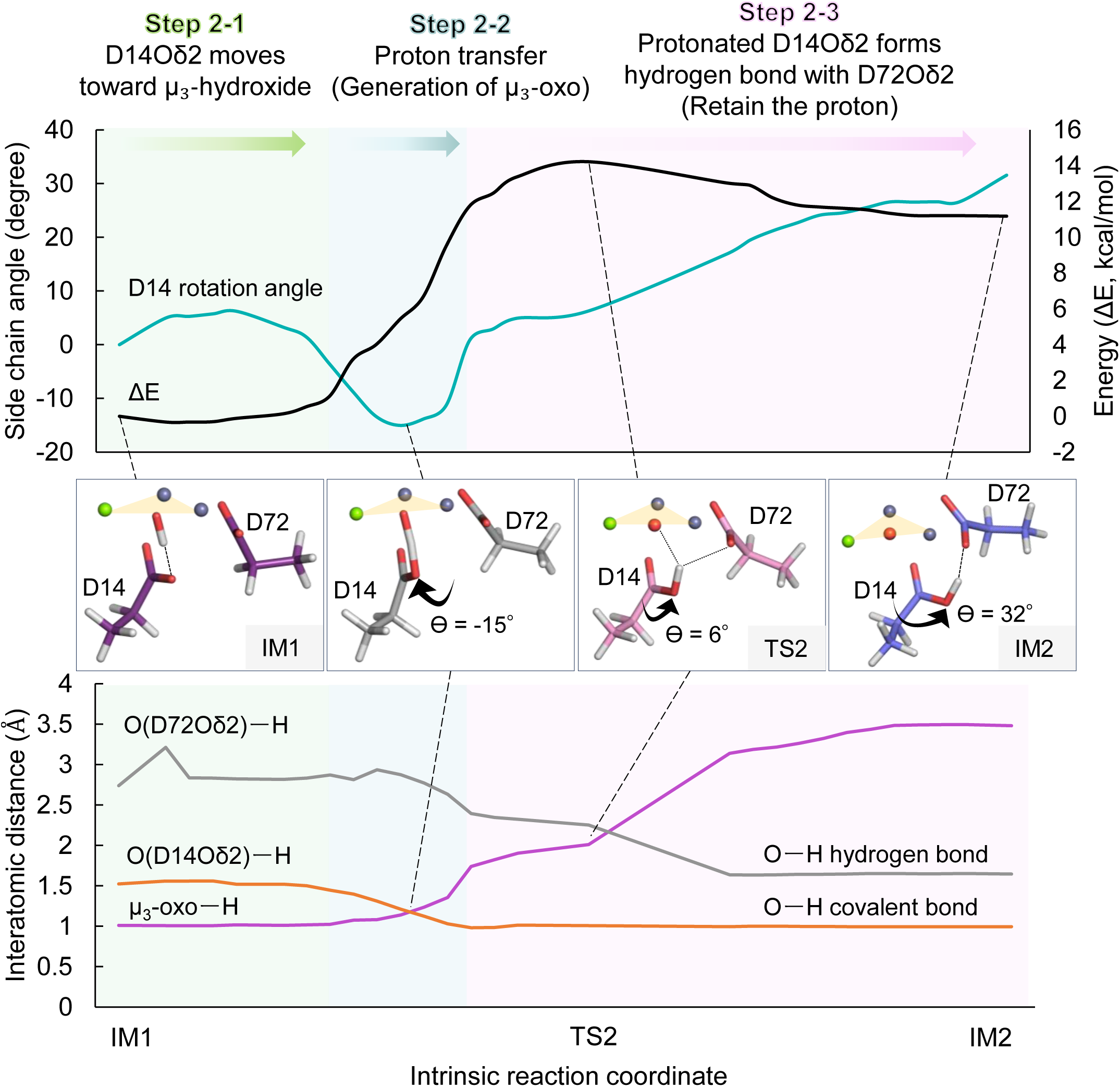
Formation of the µ₃-oxo species in the POP-bound Zn²⁺–ShPPase. Upper graph: rotation angle of Asp14 (light blue) and energy profile (black) during the transition from IM1 to IM2. Lower graph: interatomic distances: O(D72Oδ2)–H (grey), Asp14Oδ2–H (orange), and µ₃-oxo–H (pink). Three mechanistic steps—Step 2-1: Asp14Oδ2 approaches µ₃-OH (green), Step 2-2: proton transfer (blue), Step 2-3: protonated Asp14 forms H-bond with Asp72 (pink)— are highlighted. Optimized key intermediates and transition state structures are shown as bold stick models in black boxes. Full active-site models are shown in Fig. S6. Atom colouring is as in Fig. 3.

In step 2-1, the Oδ2 atom of Asp14 approaches the proton of the μ₃-hydroxide, positioning it between the two oxygen atoms. As negative rotation proceeds (step 2-2), the proton is transferred to Oδ2, forming a covalent O–H bond and generating the μ₃-oxo species. The crossing of the Asp14 Oδ2–H and μ₃-oxo–H distances in this step reflects proton transfer (Fig. 5). Although this step raises the system’s energy, the true transition state occurs later. Comparison of partial charges for the transferring proton in enzyme and non-enzyme models showed greater polarization by 0.436 in Mulliken charge in the enzymatic context, suggesting that the trinuclear metal center facilitates deprotonation by lowering the p*K*_a_ of the hydroxide. In step 2-3, the protonated Asp14 rotates further toward Asp72 and forms a stabilizing hydrogen bond with its side chain (Fig. 5). The rotation of protonated Asp14 corresponds to the rate-limiting step of the reaction. The point where the μ₃-oxo–H and Asp72 Oδ2–H distances cross coincides with the computed transition state, indicating that deprotonation completes as the hydrogen bond acceptor shifts from μ₃-oxo to Asp72.

Stage 3: S_N_2-type hydrolysis by the μ₃-oxo nucleophile.

In the final stage, the strongly nucleophilic μ₃-oxo species attacks the phosphorus atom (P1) of POP via TS3, forming the product complex with a low activation barrier of 2.2 kcal/mol (Figs. 3, 6a). In the IM2 state, the deprotonated μ₃-oxo is slightly offset from the Zn–Zn–Mg triangular plane and positioned 3.04 Å from P1 (Fig. 6a). At the TS, the μ₃-oxo aligns at the center of the trinuclear metal cluster, establishing an ideal geometry for in-line S_N_2 attack. As the reaction proceeds from IM2 to the product state, the O–P1–μ₃-oxo angle shifts from 164.5° to 169.1°, approaching the optimal 180° alignment^30^ supported by surrounding residues and metal ions.

**Fig. 6.**
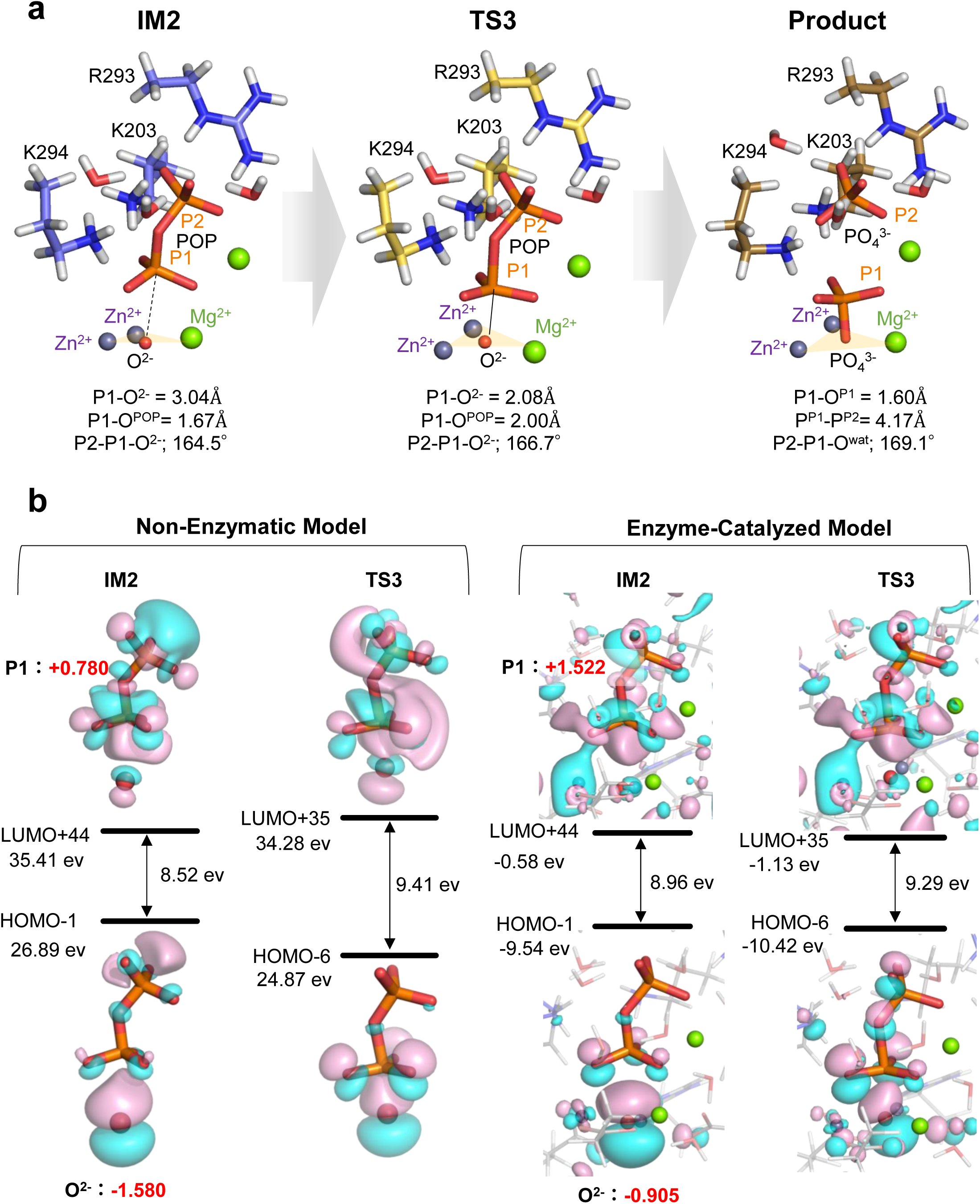
S_N_2 hydrolysis mechanism mediated by the µ₃-oxo species. **a**, Optimized structures of IM2 (blue), TS3 (yellow), and product state (brown), with key interatomic distances and angles shown. **b**, Molecular orbital maps comparing the non-enzymatic and enzyme-catalysed models. Red numbers indicate Mulliken atomic charges. Only the catalytic core is shown; for the full active site, see Fig. S6. Atom colouring follows Fig. 3.

DFT calculations delineated a plausible reaction path involving intermediates (IM) and transition states (TS) (Fig. 3). Deprotonation of the μ₃-hydroxide proceeds via TS2 with a modest activation barrier of 13.1 kcal/mol, and the fully formed μ₃-oxo appears in IM2. The overall activation energy of the entire reaction is 15.5 kcal/mol, consistent with efficient catalysis at physiological temperature (∼300 K). These findings indicate that Asp14-mediated deprotonation of the μ₃-hydroxide is the rate-limiting step of the hydrolysis reaction, and that the formation of the μ₃-oxo species is key to enhancing nucleophilic reactivity.

Electrostatic potential (ESP) maps of IM1 and IM2 show the μ₃-oxo carrying a highly negative potential (red region), confirming its strong nucleophilic character (Fig. S11). In contrast, the μ₃-hydroxide exhibits lower nucleophilicity (Fig. S11). Mulliken charge analysis supports this, with the μ₃-oxo carrying a charge of –0.91 and the μ₃-hydroxide –0.83. These results provide a mechanistic rationale for the higher hydrolytic activity of Family II PPases relative to their Family I counterparts, which lack the ability to generate a highly nucleophilic μ₃-oxo intermediate.

### Catalytic role of the trinuclear metal center in S_N_2-type hydrolysis by Family II PPase

As demonstrated above, the μ₃-oxo species in Family II PPase functions as a highly nucleophilic intermediate and is central to catalytic activity. In this section, we further dissect the electronic and mechanistic properties of this intermediate using molecular orbital (MO) theory–based calculations. One of the principal functions of the trinuclear metal center is to lower the p*K*a of the μ₃-hydroxide oxygen, thereby facilitating its deprotonation and enabling the formation of a strongly nucleophilic oxo species. This structural arrangement not only promotes the generation of a reactive intermediate but also enhances the overall efficiency of the S_N_2-type hydrolysis via multiple catalytic strategies.

MO analysis revealed that the electron density of the oxo ion (O²⁻) is more symmetrically distributed in the enzyme-catalyzed system compared to the non-enzymatic model, indicating stabilization of this inherently reactive species (Fig. 6b). Moreover, the trinuclear metal center positions the oxo nucleophile in an optimal geometry for attack on the phosphorus atom of POP, while simultaneously coordinating the three oxygen atoms of the substrate. In addition, the enzyme active site facilitates charge redistribution at the reaction center. Mulliken charge analysis showed that the phosphorus atom (P1) carries a charge of +1.522 in the enzymatic system versus +0.780 in the non-enzymatic system, indicating that coordination to the metal ions enhances the electrophilicity of the phosphorus by drawing electron density away.

During the transition from IM2 to TS3, the total charge of the three oxygen atoms bonded to P1 changes significantly in the non-enzymatic model (–0.146), but remains nearly constant in the enzymatic model (+0.006), clearly showing that metal–oxygen interactions stabilize the transition state.

These results collectively demonstrate that the trinuclear metal center of Family II PPase integrates multiple catalytic functions to facilitate highly efficient hydrolysis. In particular, deprotonation of the μ₃-hydroxide by Asp14 yields a highly reactive oxo species that is stabilized within the trinuclear environment. This arrangement ensures an ideal in-line configuration of the nucleophile and leaving group, thereby significantly lowering the activation energy compared to non-enzymatic systems.

These insights provide a general mechanistic framework for metal-dependent cleavage of P–O bonds, particularly among enzymes classified as type E phosphatases. They also offer valuable guidance for the rational design of metal-based catalysts and artificial metalloenzymes. Furthermore, these findings may inform the development of metalloprotein-inspired therapeutics and biomimetic protein engineering strategies.

### Understanding the formation of active species

Metalloproteins utilize a variety of highly evolved strategies to generate highly reactive transition metal species—such as metal–oxo/oxyl, metal–superoxo, and metal–(hydro)peroxo intermediates—that enable extremely efficient catalysis by significantly lowering activation energies^31^. However, the mechanisms underlying the formation of these active species remain poorly understood. For example, although a μ₃-oxo species has been proposed at the trinuclear metal site of PhoX-type phosphatases involved in P–O bond cleavage, the specific formation pathway has yet to be elucidated^6,17^.

In this study, we focused on the early activation step—i.e., the “induction phase”—to uncover the mechanism underlying the formation of the active nucleophile. Our results revealed that Asp14 acts as a general base, and that its unique conformational rotation plays a decisive role in facilitating deprotonation (Figs. 3–5). Structural analysis showed sufficient spatial flexibility around Asp14 to allow this rotation. In addition, site-directed mutagenesis of Asp14, combined with activity assays and sequence conservation analysis, confirmed its essential catalytic function (Table 1, Figs. S8, S10). These findings suggest that the precise spatial positioning of amino acid residues to act as catalytic bases represents a sophisticated and evolutionarily optimized mechanism for controlling active species formation in metalloproteins.

By integrating experimental XCS-EXAFS data with density functional theory (DFT) calculations, we have uncovered a complete mechanistic model for Family II PPase catalysis, including the crucial activation phase. Notably, we found that the energy barrier associated with μ₃-oxo formation is higher than that of subsequent catalytic steps, indicating that the generation of the reactive intermediate constitutes the rate-limiting step of the entire reaction. This implies that active species formation, rather than substrate transformation per se, is the major determinant of reaction efficiency.

While many studies have focused on the hydrolysis pathway or product formation, the initial step of how the active species forms has often been overlooked. By highlighting this underappreciated yet fundamental step, our findings suggest new directions for enhancing catalytic performance and optimizing artificial enzymes and metal complexes. We propose that future efforts to understand and engineer metalloenzymes should place greater emphasis on optimizing the formation of active species.

## Conclusions

In this study, we elucidated the catalytic mechanism of Family II inorganic pyrophosphatase (PPase) using a combination of X-ray crystallography-based extended X-ray absorption fine structure (XCS-EXAFS) analysis, mutagenesis experiments, and density functional theory (DFT) calculations. We identified a key reaction pathway involving the deprotonation of a μ₃-hydroxide intermediate by the conserved residue Asp14, followed by the formation of a highly nucleophilic μ₃-oxo species that initiates S_N_2-type hydrolysis of inorganic pyrophosphate. Quantum chemical analyses revealed that the trinuclear metal center facilitates oxo formation by lowering the p*K*a of the hydroxide species and stabilizing the reactive intermediate, aligns the μ₃-oxo species for optimal nucleophilic attack, and stabilizes the transition state. These findings establish the formation of the μ₃-oxo species as the rate-limiting step of catalysis and highlight the critical role of active species generation in determining reaction efficiency. Our study provides mechanistic insights into metal-dependent phosphate hydrolysis and offers a framework for the rational design of artificial metalloenzymes and bioinspired catalysts.

## Methods

### Materials and Protein Purifications

The recombinant wild-type ShPPase and ShPPase variants were expressed in *E. coli* BL21 (DE3) using pET16b as an expression vector and purified as previously described^32^. Metal-free ShPPase was prepared by EDTA treatment. The enzyme was diluted with 100 mM Tris/HCl (pH 7.5) buffer containing 20 µM EDTA and 50 mM KCl, subjected to three dilution/concentration cycles using ultrafiltration.

### Sample preparation for EXAFS

For EXAFS studies, 0.5 mg/mL metal-free ShPPase samples were incubated in an activation buffer containing 100 mM Tris/HCl (pH 7.5), 20 µM EDTA, 50 mM KCl, and 0.5 mM ZnCl_2_ for two hours at 4°C. Samples were desalted using spin columns two or three times to eliminate excess metals, which cause noise in EXAFS studies. During the spin column process, the metal-free buffer with 40 µM ZnCl_2_ maintains two combined metals in the active site. This low concentration effect of ZnCl_2_ can be ignored for EXAFS studies because it accounts for only 0.0004 % of the total sample. Substrate-free Zn^2+^-ShPPase sample was loaded into polypropylene PCR tubes, flash-frozen in liquid-nitrogen, and stored in a −80 °C freezer. PNP-bound Zn^2+^-ShPPase sample contained final concentrations of 1.1 mM Zn^2+^-ShPPase, 0.1 M PNP, and 0.1 M MgCl_2_ at pH 7.5, and was stored in the same way as the substrate-free Zn^2+^-ShPPase sample. Zn^2+^-ShPPase samples had sufficient activity without extra metals (Fig. S8, Table S1).

### X-ray absorption spectroscopy

X-ray absorption fine structure (XAFS) is a structural technique that probes the local environment of a metal ion at a 0.01 Å-level resolution. For XAFS data collected at the metal *K*-edge, the edge region of the spectrum is sensitive to the effective charge on the metal ion and the coordination geometry, while the EXAFS region provides average metal–ligand bond distances and coordination numbers.

X-ray absorption measurements of each sample were obtained at the beamline 15 of the SAGA Light Source (SAGA-LS), with the ring operating at 1.4 GeV, 100-300 mA. A Si (111) double-crystal monochromator was used for energy selection at the Zn K-edge and data were measured in fluorescence mode as Zn Kα fluorescence using a 7-element Silicon drift detector (SDD, Techno-AP). The position of the SDD was adjusted to suppress the dead time to under 5%. All Zn XAFS spectra were collected by scanning the incident X-ray energy from 9.3310 to 10.2096 keV. The averaged data included 10 scans at about 42 minutes per scan for each sample to increase the signal-to-noise ratio. The sample was maintained under the supercooled environment in a stream of 100 K-nitrogen gas during data collection to minimize radiation damage. No photodegradation was observed for any of the samples from XANES data and activity assay (Fig. S8, Table S2). In all experiments, individual scans were normalized to the incident photon flux and averaged using the program Athena from the software package Demeter^23^. For the averaged data, further processing of spectra including background subtraction and normalization was also performed using Athena, following standard protocols for X-ray spectroscopy described below. The data were normalized to an edge jump of 1.0 between background and spline curves at 9660 eV.

EXAFS fitting was performed using the program Artemis, also of the software package Demeter^23^. Possible scattering paths for the EXAFS models were initially determined using FEFF 7.0 in combination with a recent high-resolution crystal structure (PDB ID: 6ll7,6ll8).

Each fit used a common value *Δ*E_0_ for every component in the fit. With a value of S_0_^2^, the scale factor was restricted to 0.8-1.0 for reproducibility. Experimental EXAFS data were converted to *k* space by applying the EXAFS equation and subsequently weighted by *k*^3^ to compensate for the damping of oscillations at high *k*. The *k*^3^ data were fit over similar *k*-ranges (*k*=3 to 9) using a nonlinear least-squares approach with theoretical values for Zn–O, Zn–N, Zn–P, and Zn–Zn bonds from FEFF 7.0 using structural refinement data for Zn^2+^-ShPPase. EXAFS spectra were also Fourier transformed to produce radial structure functions (RSFs) that isolate frequency correlations between the central absorbing atom (Zn) and neighboring atoms as a function of bond distance (*R*). All data were fit in *R*-space using an *R*-range of 1 to 3.5 Å. Due to the complexity of the EXAFS of the ShPPase, fitting was limited to include only single scattering paths. No smoothing was used at any point in any of the data processing.

### Activity assay and kinetic analysis

The activity was measured by the molybdenum blue method. A reaction mixture containing 10 μL of enzyme and 110 μL of 1 mM potassium pyrophosphate (K_4_POP) as substrate in 100 mM Tris-HCl buffer, 50 mM KCl (pH 7.5), containing 5 mM MgCl_2_, was incubated for 3 min at 25 °C. The reaction was stopped by the addition of 30 μL of 50 mM H_2_SO_4_. The reaction mixture was colored by the addition of 150 μL of 1 % ammonium molybdate in 0.05 % K_2_SO_4_ and 1 % sodium ascorbate in Milli-Q water. The amount of phosphate liberated from the hydrolysis of inorganic POP was measured at 750 nm using a microplate reader (Thermo Fisher Scientific, Waltham, MA, USA) and a standard phosphate curve (0-500 μM phosphate) after 20 minutes. Specific activity (U/mg) was defined as 1 μmol of POP hydrolyzed per min per mg of protein. One unit of activity corresponded to the formation of 2 µmol of phosphate per min from 1 µmol of POP under the assay conditions.

### DFT calculations

First, the X-ray crystal structure (XCS) model was constructed based on the Mg²⁺-ShPPase– PNP complex (PDB ID: 6ll8). Building upon this structural model, two additional computational models were constructed: an extended X-ray absorption fine structure (EXAFS) simulation model and a proposed hydrolysis reaction mechanism model (POP). The XCS model included the PNP substrate analogue, four Mg atoms, the F atom as a center of tri-metal structure, 11 water molecules, and the side chains of His8, Asp12, Asp14, Asp72, Asp93, His94, His95, Asp146, Lys203, Arg293, and Lys294. The total number of atoms and the total charge are 187 and +1, respectively. In the EXAFS model, two Mg atoms as binuclear metals were replaced by two Zn atoms. The F atom as a center of tri-metal structure was replaced by the O atom or OH molecule. The total number of atoms and the total charge are 188 and +1, respectively. In the POP model, the N atom of PNP was replaced with the O atom, the F atom as a center of tri-metal structure was replaced by the O atom or OH molecule, and two Mg atoms as binuclear metals were replaced by two Zn atoms. The total number of atoms and the total charge are 188 and +2, respectively. We also constructed the Asp14 rotation model for Asp14 rotation hypothesis, which added three side chains of Cys115, Ser116, and Asn117 to the POP model. The total number of atoms and the total charge are 222 and +2, respectively. In all calculated models, most of the side chains of amino acid residues were truncated at their *β*-carbon, which were converted to methyl groups. During geometry optimization, the *β*-carbon atoms were fixed to their crystallographic positions. For Lys203, Arg293, and Lys294 in each model, the frozen atom was selected to be the *γ*-carbon to limit the size of the Quantum Mechanics (QM) region to mimic the enzyme environment and prevent the unrealistic distortion of the cluster and ensure sufficient flexibility. The QM subsystem was treated using density function theory (DFT) calculations. All DFT calculation was performed using the B3LYP functional^33,34^ and the 6-31G(d) basis set^35,36^ for H, C, N, O, P, S, F, and 6-311+G(d) basis set^37,38^ for Mg, Zn, implemented with the Gaussian 16 program package.^39^ The protonated model was characterized by the Amber force field (Amber10:EHT)^40^ using the Molecular Operating Environment (MOE).^40^ Vibrational frequency analyses were performed at the same level of theory to confirm that the optimized structures correspond to true minima (no imaginary frequencies) or transition states (one imaginary frequency). However, the imaginary freqencies due to the constrained carbon atoms were ignored. All thermodynamic parameters and Gibbs energy corrections were obtained from these frequency calculations at 298.15 K. The reaction pathway was traced by the quasi-intrinsic reaction coordinate (quasi-IRC) method, followed to obtain the minimum-energy path, in which the TS structure was slightly perturbed in the direction of the reactants and products, followed by full geometry optimizations except the constrained carbon atoms.

## Data Availability

All data pertaining to this study have been presented in the manuscript.

## Acknowledgments

This work was supported by a Grant-in-Aid for Scientific Research from the Ministry of Education, Culture, Sports, Science and Technology, Japan (JSPS KAKENHI; grant nos. JP23KJ1741 (S.M.), JP23K26836 (M.H.), JP24K21245 (K.Y.) and JP25K08678 (Y.S.)). We thank the Center for Advanced Instrumental and Educational Support of the Faculty of Agriculture (Kyushu University) for use of facilities for the enzyme assay.

## Author contributions

S.M. and K.W. conceived the study and designed the research. S.M. and M.H. prepared the protein samples. S.M., E.M., H.S., M.K., M.H. and K.W. performed the XAFS experiments, and S.M., E.M., H.S. and K.W. analysed the data. S.M., T.T. and Y.Kak. conducted the mutagenesis experiments. S.M., Y.S., K.Y. and K.W. designed the DFT calculations, S.M. and Y.Kam. carried out the calculations, and S.M., Y.Kam., Y.S., K.Y., T.T., Y.Kak. and K.W. discussed the results. S.M. and K.W. wrote the paper with input from all authors.

## References

1. Chalkley, M. J., Mann, S. I. & DeGrado, W. F. De novo metalloprotein design. Nat. Rev. Chem. 6, 31–50 (2021).

2. Zastrow, M. L. & Pecoraro, V. L. Designing functional metalloproteins: from structural to catalytic metal sites. Coord. Chem. Rev. 257, 2565–2588 (2013).

3. Maret, W. Metalloproteomics, metalloproteomes, and the annotation of metalloproteins. Metallomics 2, 117–125 (2010).

4. Westheimer, F. H. Why nature chose phosphates. Science 235, 1173–1178 (1987).

5. Wagner, C. A. The basics of phosphate metabolism. Nephrol. Dial. Transplant. 39, 190– 201 (2024).

6. Ji, J.-N. & Chen, S.-L. μ_3_-Oxo stabilized by three metal cations is a sufficient nucleophile for enzymatic hydrolysis of phosphate monoesters. Dalton Trans. 45, 2517–2522 (2016).

7. Tonks, N. K. Protein tyrosine phosphatases: from genes, to function, to disease. Nat. Rev. Mol. Cell Biol. 7, 833–846 (2006).

8. Zhang, Z.-Y. Protein tyrosine phosphatases: structure and function, substrate specificity, and inhibitor development. Annu. Rev. Pharmacol. Toxicol. 42, 209–234 (2002).

9. O’Brien, P. J. & Herschlag, D. Alkaline phosphatase revisited: hydrolysis of alkyl phosphates. Biochemistry 41, 3207–3225 (2002).

10. Bobyr, E. et al. High-resolution analysis of Zn^2+^ coordination in the alkaline phosphatase superfamily by EXAFS and X-ray crystallography. J. Mol. Biol. 415, 102–117 (2012).

11. Guddat, L. W. et al. Crystal structure of mammalian purple acid phosphatase. Structure 7, 757–767 (1999).

12. Schenk, G., Mitić, N., Hanson, G. R. & Comba, P. Purple acid phosphatase: a journey into the function and mechanism of a colorful enzyme. Coord. Chem. Rev. 257, 473–482 (2013).

13. Shevelev, I. V. & Hübscher, U. The 3′–5′ exonucleases. Nat. Rev. Mol. Cell Biol. 3, 364– 376 (2002).

14. Aubert, S. D., Li, Y. & Raushel, F. M. Mechanism for the hydrolysis of organophosphates by the bacterial phosphotriesterase. Biochemistry 43, 5707–5715 (2004).

15. Chen, S.-L., Fang, W.-H. & Himo, F. Theoretical study of the phosphotriesterase reaction mechanism. J. Phys. Chem. B 111, 1253–1255 (2007).

16. Ariño, J., Velázquez, D. & Casamayor, A. Ser/Thr protein phosphatases in fungi: structure, regulation and function. Microb. Cell 6, 217–256 (2019).

17. Yong, S. C. et al. A complex iron-calcium cofactor catalyzing phosphotransfer chemistry. Science 345, 1170–1173 (2014).

18. Kajander, T., Kellosalo, J. & Goldman, A. Inorganic pyrophosphatases: one substrate, three mechanisms. FEBS Lett. 587, 1863–1869 (2013).

19. Terkeltaub, R. A. Inorganic pyrophosphate generation and disposition in pathophysiology. Am. J. Physiol. Cell Physiol. 281, C1–C11 (2001).

20. Horitani, M. et al. X-ray crystallography and electron paramagnetic resonance spectroscopy reveal active site rearrangement of cold-adapted inorganic pyrophosphatase. Sci. Rep. 10, 4368 (2020).

21. Fabrichniy, I. P. et al. A trimetal site and substrate distortion in a Family II inorganic pyrophosphatase. J. Biol. Chem. 282, 1422–1431 (2007).

22. Baykov, A. A., Anashkin, V. A., Salminen, A. & Lahti, R. Inorganic pyrophosphatases of Family II —two decades after their discovery. FEBS Lett. 591, 3225–3234 (2017).

23. Ravel, B. & Newville, M. ATHENA and ARTEMIS interactive graphical data analysisusing IFEFFIT. Physica Scripta 1007 (2005)

24. Riggs-Gelasco, P. J., Stemmler, T. L. & Penner-Hahn, J. E. XAFS of dinuclear metal sites in proteins and model compounds. Coord. Chem. Rev. 144, 245–286 (1995).

25. Hendrickson, W. A., Co, M. S., Smith, J. L., Hodgson, K. O. & Klippenstein, G. L. X-ray absorption spectroscopy of the dimeric iron site in azidomethemerythrin from Phascolopsis gouldii. Proc. Natl. Acad. Sci. U.S.A. 79, 6255–6259 (1982).

26. W. T, E. & E. A, S. An x-ray absorption study of the binuclear iron center in deoxyhemerythrin. J. Am. Chem. Soc. 7, 1919–1923 (1983).

27. Ginting, E. L., Maeganeku, C., Motoshima, H. & Watanabe, K. Functional characteristics of inorganic pyrophosphatase from psychrotroph Shewanella sp. AS-11 upon activation by various divalent cations. Asian J. Chem. 26, 611–616 (2014).

28. Fabrichniy, I. P. et al. Structural studies of metal ions in Family II Pyrophosphatases: the requirement for a janus ion,. Biochemistry 43, 14403–14411 (2004).

29. Yang, L., Liao, R.-Z., Yu, J.-G. & Liu, R.-Z. DFT Study on the mechanism of escherichia coli inorganic pyrophosphatase. J. Phys. Chem. B 113, 6505–6510 (2009).

30. Vayner, G., Houk, K. N., Jorgensen, W. L. & Brauman, J. I. Steric retardation of S_N_2 reactions in the gas phase and solution. J. Am. Chem. Soc. 126, 9054–9058 (2004).

31. Huang, X. & Groves, J. T. Oxygen activation and radical transformations in heme proteins and metalloporphyrins. Chem. Rev. 118, 2491–2553 (2018).

32. Ginting, E. L., Iwasaki, S., Maeganeku, C., Motoshima, H. & Watanabe, K. Expression, purification, and characterization of cold-adapted inorganic pyrophosphatase from psychrophilic Shewanella sp. AS-11. Prep. Biochem. Biotechnol. 44, 480–492 (2014).

33. Becke, A. D. Density-functional thermochemistry. III. The role of exact exchange. J. Chem. Phys. 98, 5648–5652 (1993).

34. Lee, C., Yang, W. & Parr, R. G. Development of the Colle–Salvetti correlation-energy formula into a functional of the electron density. Phys. Rev. B 37, 785–789 (1988).

35. Ditchfield, R., Hehre, W. J. & Pople, J. A. Self-consistent molecular orbital methods. XII. Further extensions of Gaussian-type basis sets for use in molecular orbital studies of organic molecules. J. Chem. Phys. 56, 2257–2261 (1972).

36. Francl, M. M., et al. Self-consistent molecular orbital methods. XXIII. A polarization-type basis set for second-row elements. J. Chem. Phys. 77, 3654–3665 (1982).

37. Raghavachari, K., Binkley, J. S., Seeger, R. & Pople, J. A. Self-consistent molecular orbital methods. XX. A basis set for correlated wave functions. J. Chem. Phys. 72, 650–654 (1980).

38. McLean, A. D. & Chandler, G. S. Contracted Gaussian basis sets for molecular calculations. I. Second row atoms, Z=11–18. J. Chem. Phys. 72, 5639–5648 (1980).

39. Frisch, M. J. et al. Gaussian 16, Revision C.01 (Gaussian, Inc., Wallingford CT, 2016).

40. Molecular Operating Environment (MOE), 2024.0601 (Chemical Computing Group ULC, Montreal, QC, 2025).

