## Supplemental Data 1 for "Formation of a μ_₃_-oxo nucleophile enables efficient hydrolysis by a trinuclear metal center in Family II inorganic pyrophosphatase"

**This file includes:**

Graphical Abstract
Supplementary Figures S1–S11
Supplementary Tables S1–S2

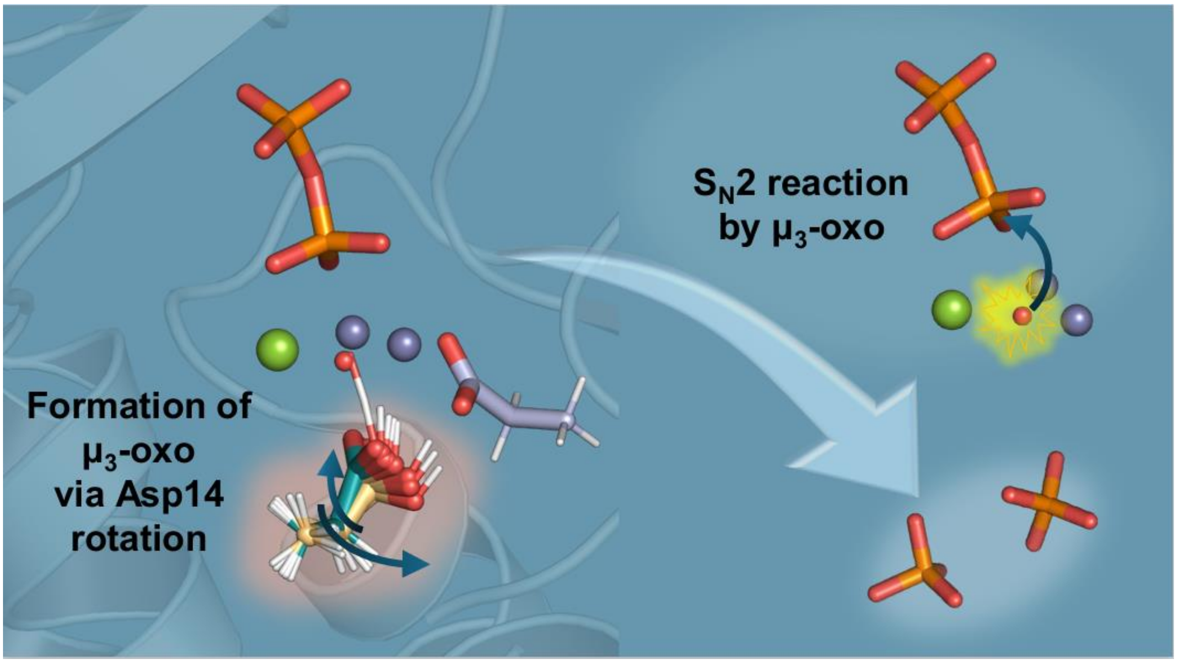


Graphical Abstract

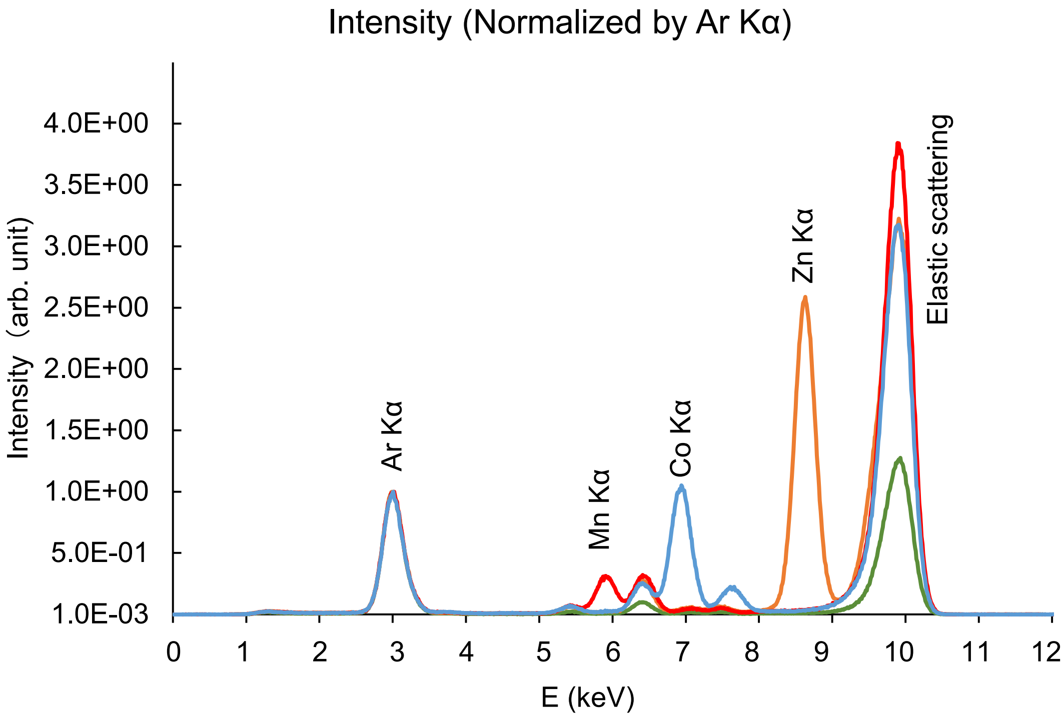


Metal-Kαemission measured by fluorescence x-ray detector. Kα emission spectra of 2 mM metal solutions as standard were measured. Zn, Mn, and Co are shown in orange, red, and light blue lines, respectively. Blank by PCR tube was shown in green line.

Fig. S1

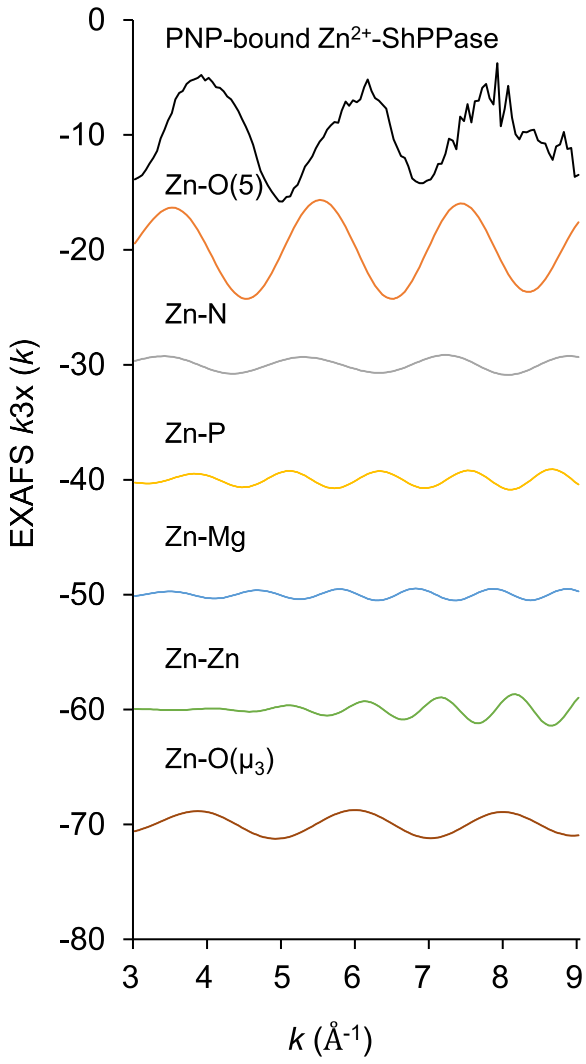


Comparison of EXAFS spectra between experimental and FEFF simulated spectra of the Zn-ShPPase. Scattering paths from FEFF include 4 oxygen atoms (orange line), a nitrogen atom (gray line), a phosphorus atom (yellow line), a magnesium atom (cyan), a zinc atom (green line), an oxygen atom derived from water which is coordinated with three metal cations (brown line)

Fig. S2

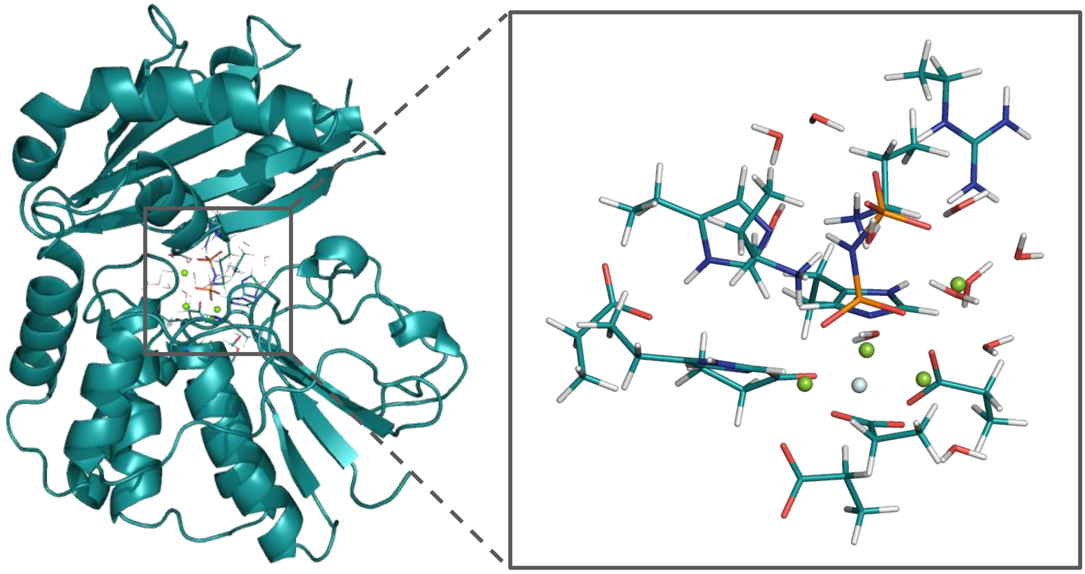

X-ray crystal structure (XSC) model of QM cluster models for the active site is depicted with the conventional licorice colors (C cyan, H white, O red, N blue, and P orange). Mg(II) ions and F are shown as green and light blue spheres, respectively.

Fig. S3

X-ray crustal structure model of PNP-bound $Mg^{2+}$-ShPPase

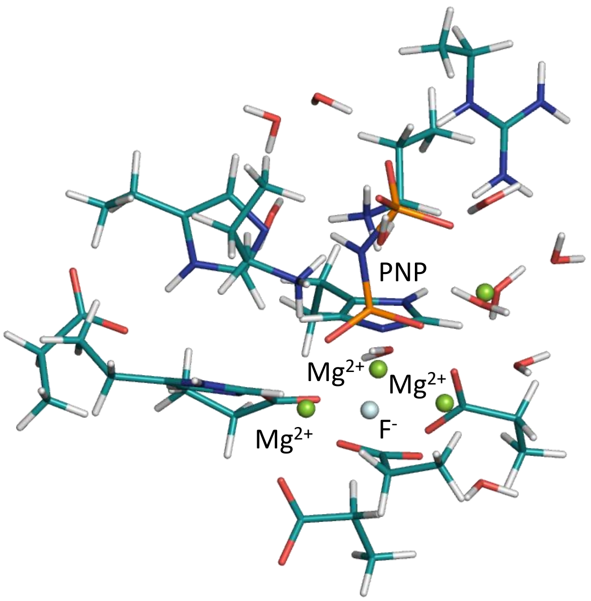


EXAFS model

**a** $\mu_3$-hydroxide state

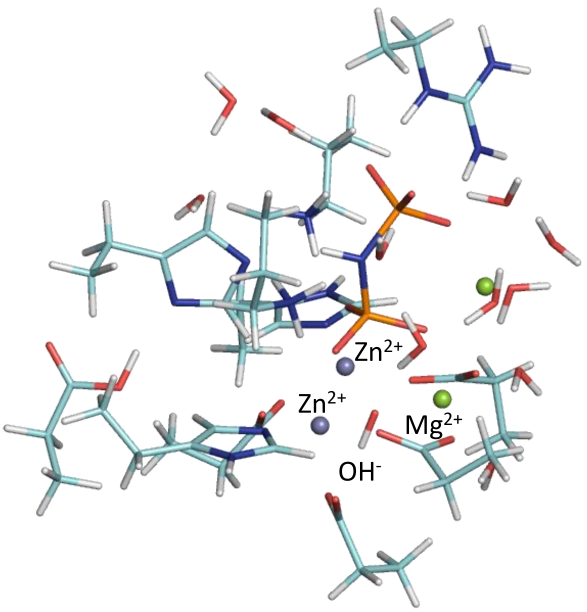


**b** $\mu_3$-oxo state

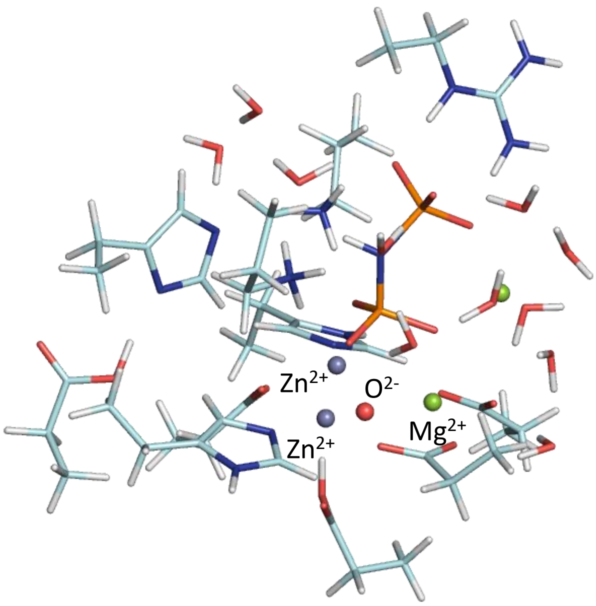


Fig. S4

Optimized structures of X-ray crystal structure models and EXAFS models ((a) $\mu_3$-hydroxide state, and (b) $\mu_3$-oxo state). The Zn, Mg, O, N, P, and H atoms are colored slate purple, green, red, blue, orange, and white, respectively.

### Asp14 rotation model with bound POP

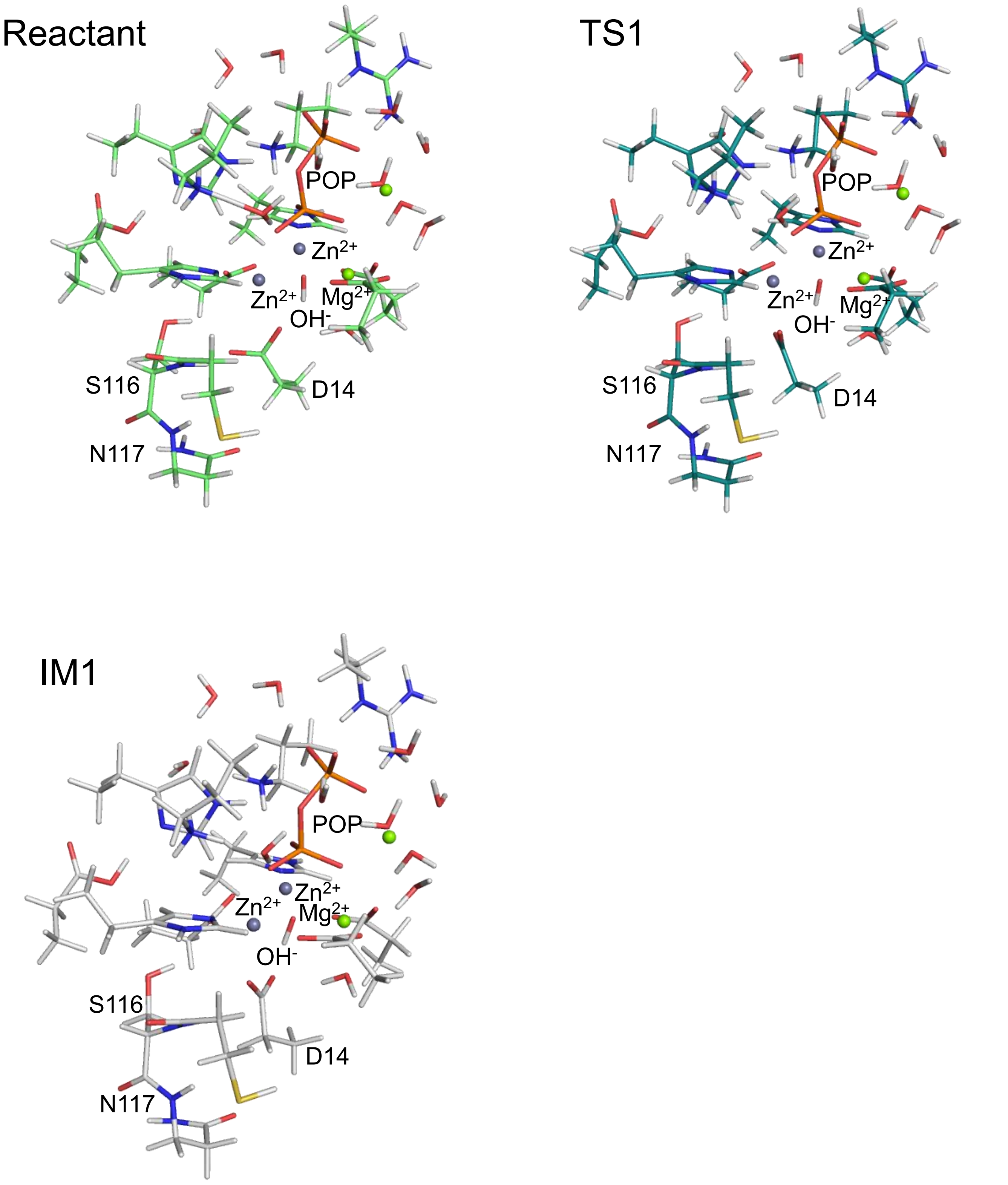


Fig. S5 Optimized structures of Asp14 rotation models. The Zn, Mg, O, N, P, and H atoms are colored slate purple, green, red, blue, orange, and white, respectively.

#### $\mu_3$-oxo formation and $S_N2$ reaction models with POP

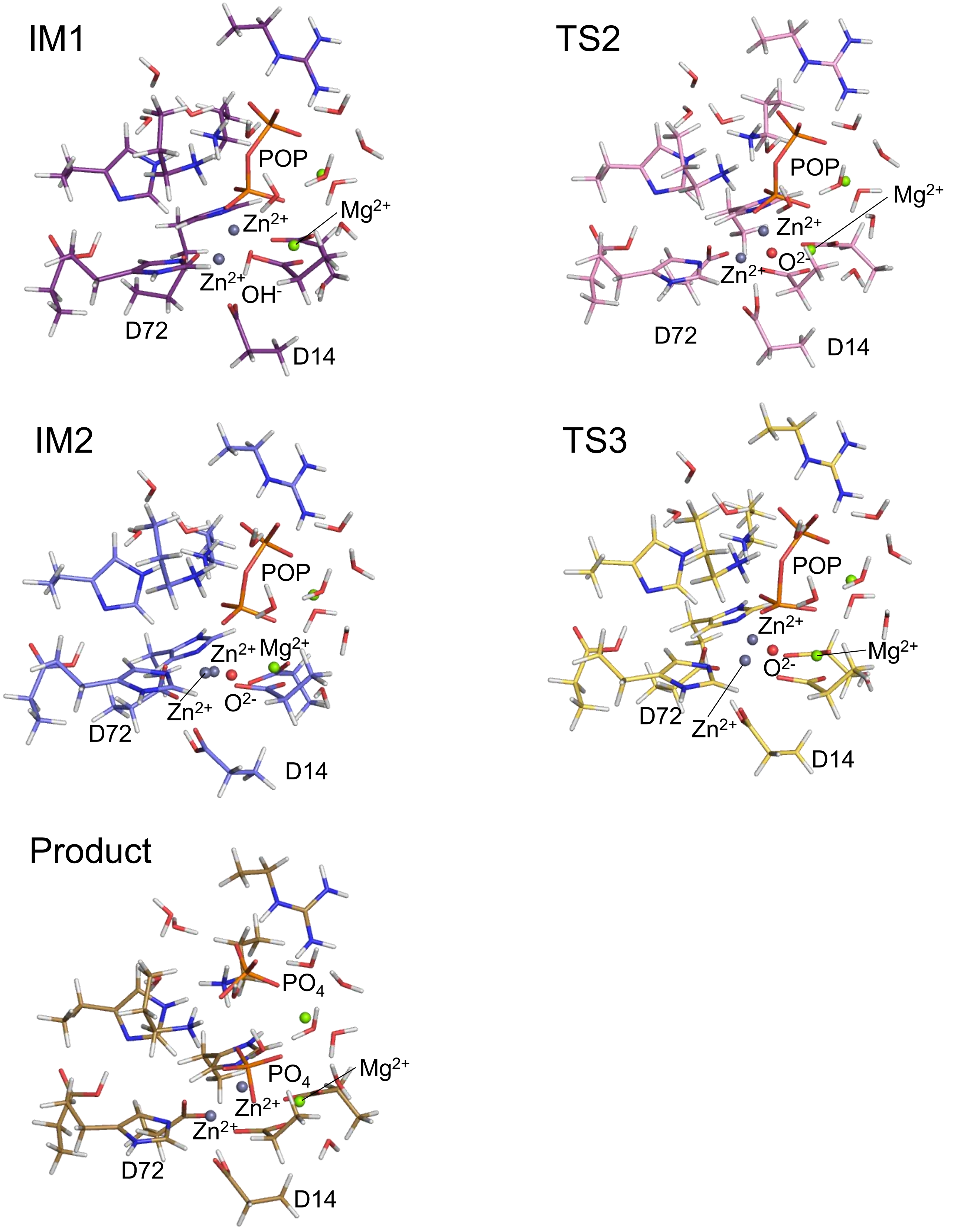


Fig. S6 Optimized structures of $\mu_3$-oxo formation and $S_N2$ reaction models. The Zn, Mg, O, N, P, and H atoms are colored slate purple, green, red, blue, orange, and white, respectively.

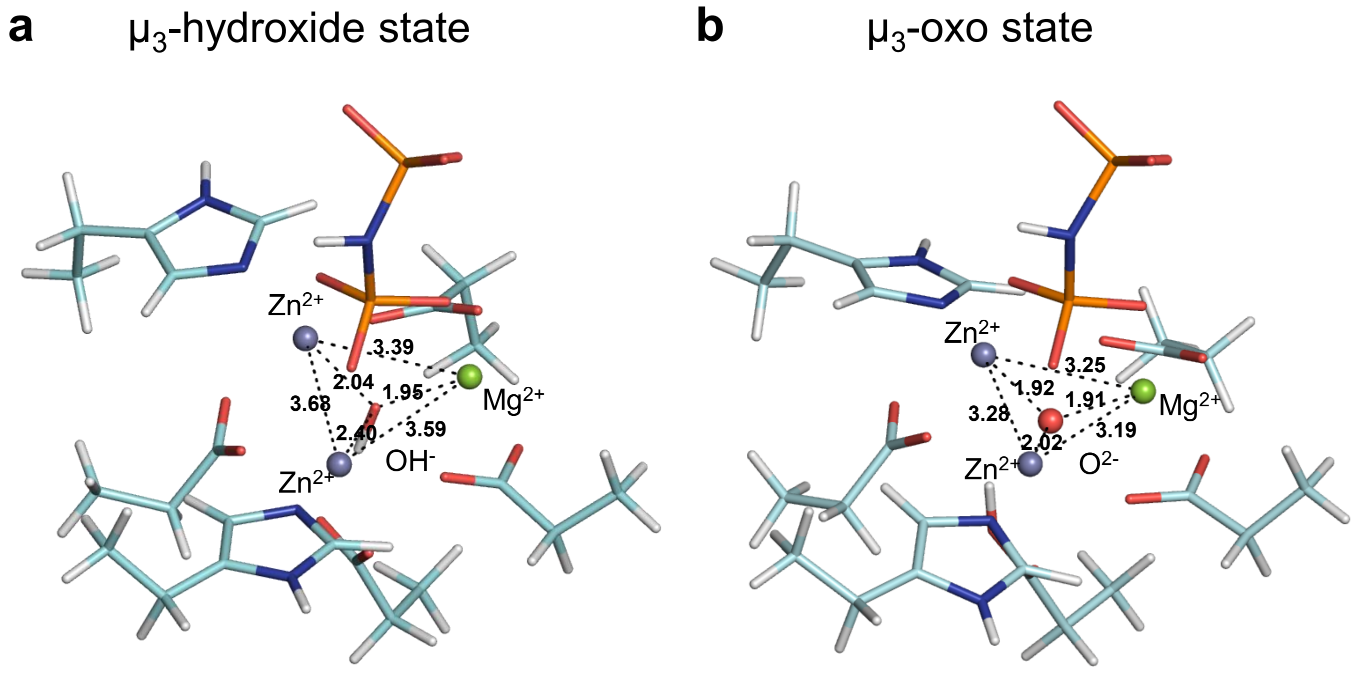


Optimized structures of EXAFS model. (a) $\mu_3$-hydroxide state, and (b) $\mu_3$-oxo state. Black dashes indicate coordination bonds. Interatomic distances are presented in Å. Only the active site near the tri-metal structure and substrate analog are shown, see Fig. S3 for EXAFS model with the whole active site. The Zn, Mg, O, N, P, and H atoms are colored slate purple, green, red, blue, orange, and white, respectively.

Fig. S7

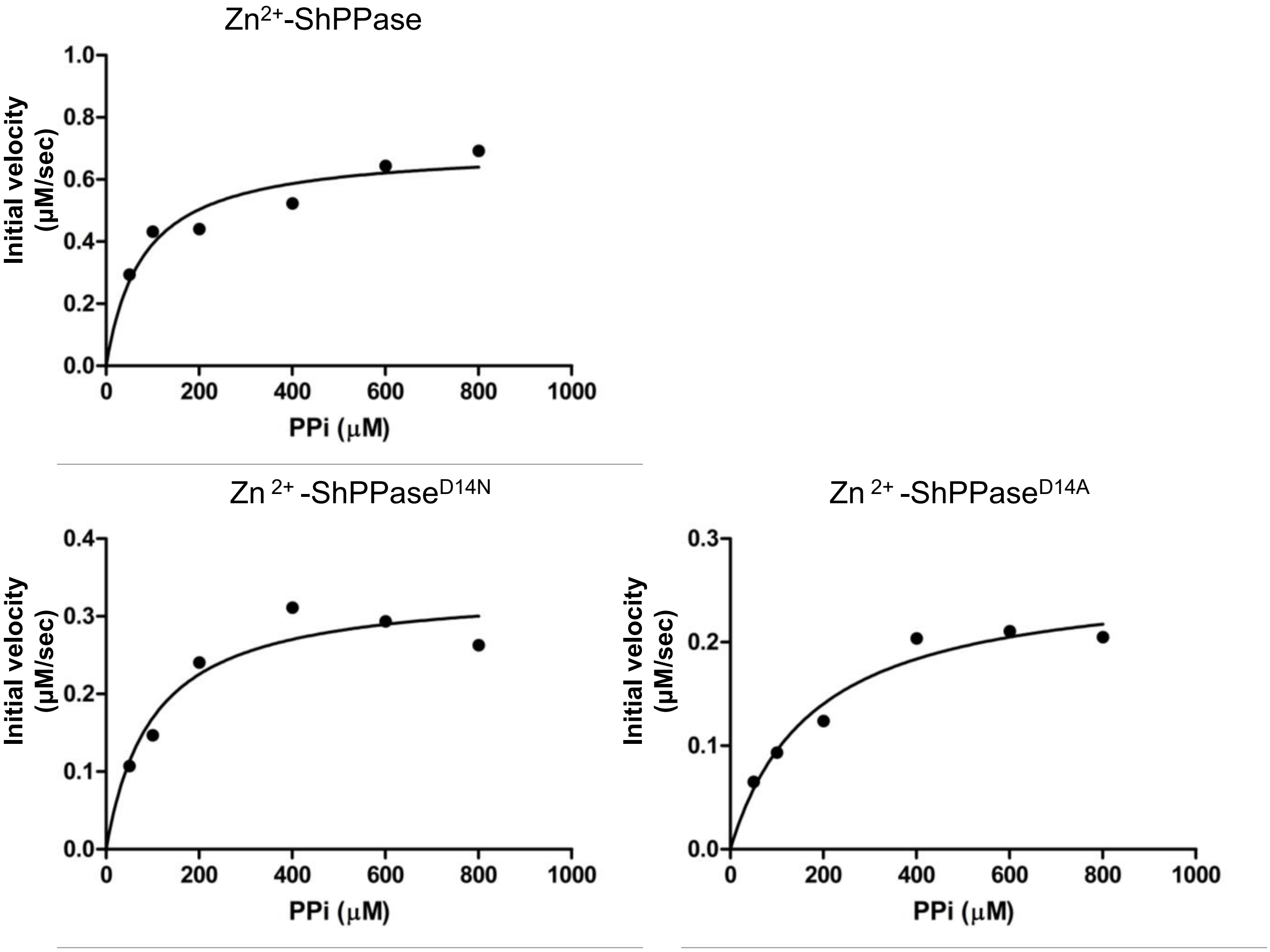


The kinetics of hydrolysis of the PPi substrate by $Zn^{2+}$-ShPPase, $Zn^{2+}$-ShPPase$^{D14N}$ and $Zn^{2+}$-ShPPase$^{D14A}$. Experimental data and fitted curves are shown in black dots and solid lines, respectively. The enzyme concentrations used in the assays were 2.9 μg/mL for $Zn^{2+}$-ShPPase, 0.17 mg/mL for $Zn^{2+}$-ShPPase$^{D14N}$, and 0.12 mg/mL for $Zn^{2}$-ShPPase$^{D14A}$.

Fig. S8

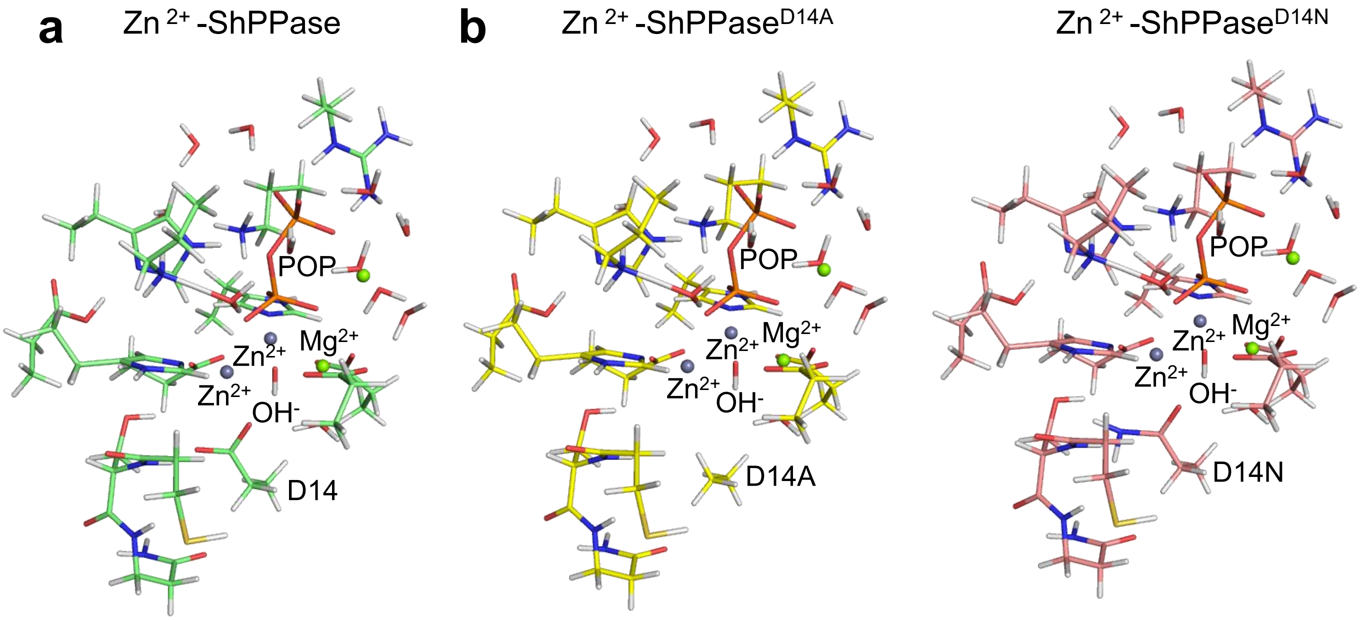


Comparison of optimized structures of (a) WT( light green) and (b) Asp14 mutation models for Zn-ShPPase$^{D14A}$ (yellow) and Zn-ShPPase$^{D14N}$ (pink). The Zn, Mg, O, N, P, and H atoms are colored slate purple, green, red, blue, orange, and white, respectively.

Fig. S9

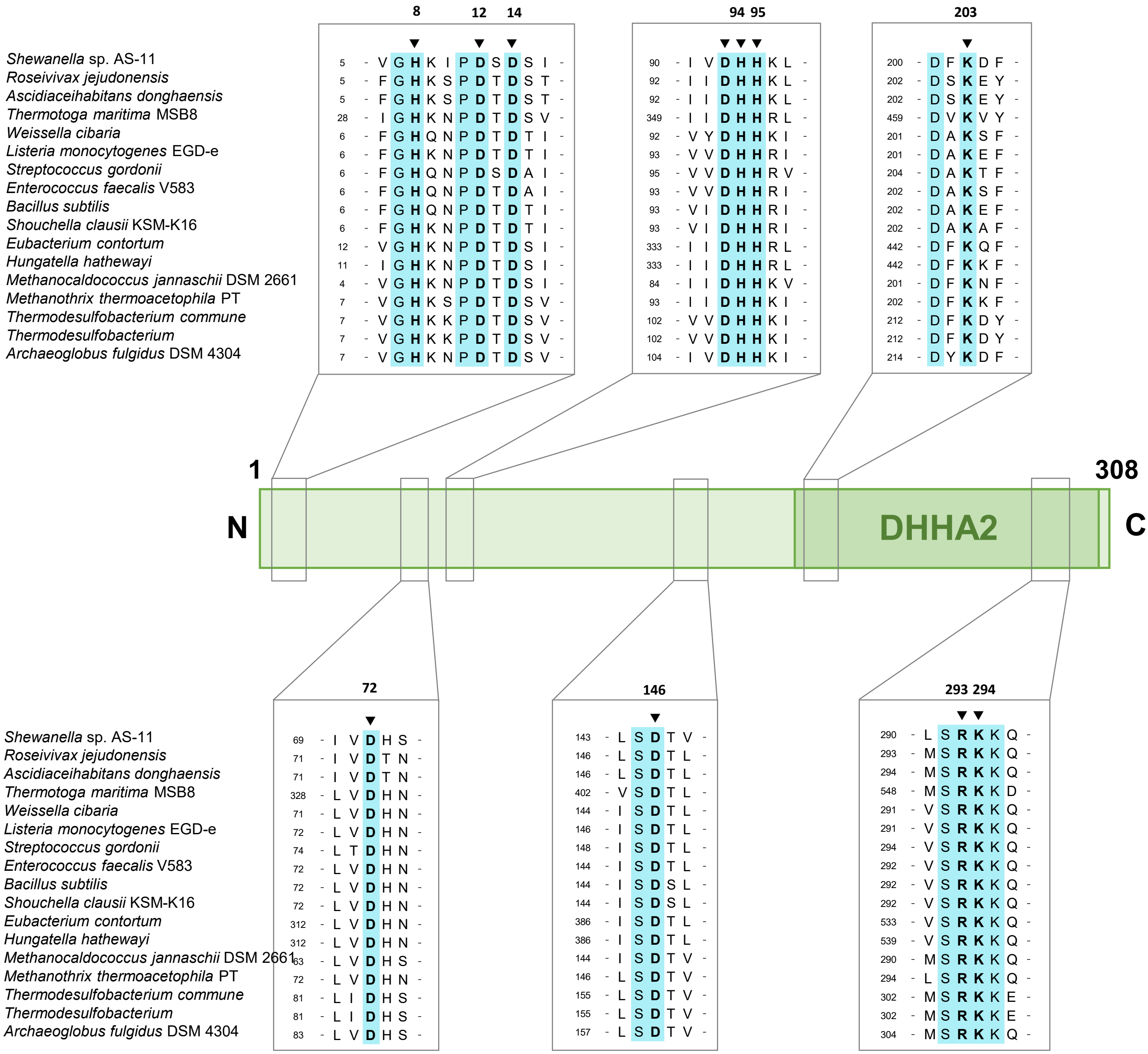


Multiple sequence alignments in reported Family II PPases. Amino acids of the active site are conserved among family II PPases and are shown in cyan (UniplotID: BAM74412.2, WP_085790359.1, SPH20124.1, AAD35672.1, KIU19850.1, CAC99526.1, AAB39104.1, AAO81394.1, BAA05186.1, BAD65437.1, CUO74905.1, CUO93176.1, AAB98601.1, ABK14034.1, HAA84199.1, WP_038063393.1, AAB90480.1). Asp 14 of ShPPase is completely conserved with others, which is shown in bold.

Fig. S10

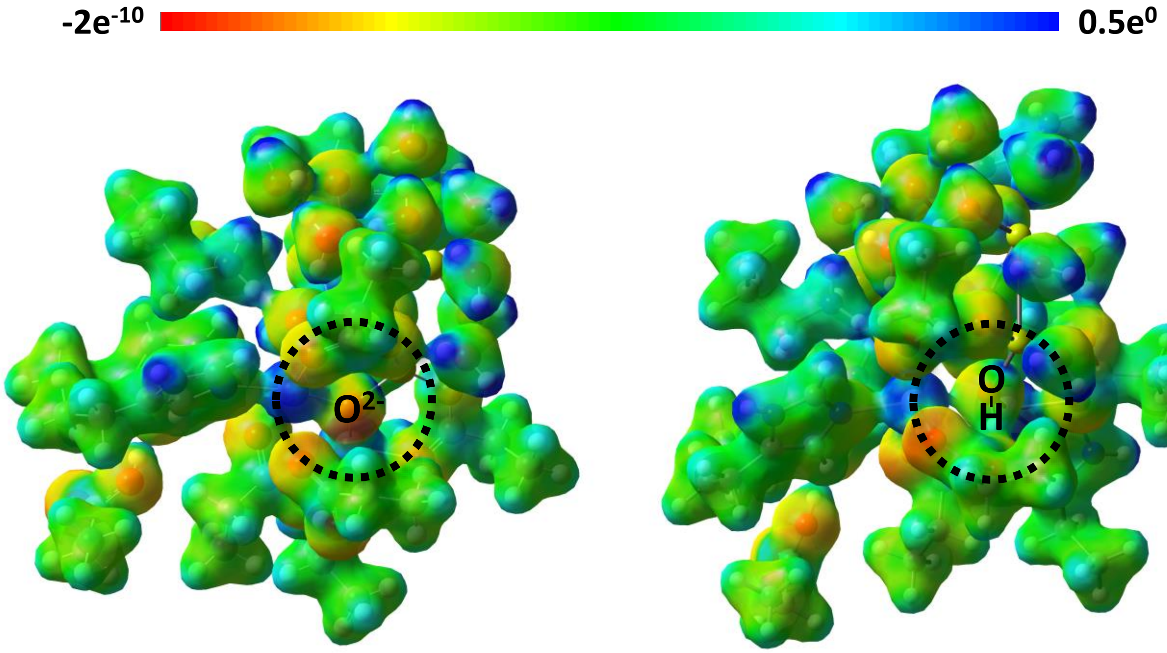


Electrostatic potential (ESP) map for IM2 state (left) and IM1 state (right). ESP of the ions of interest mapped onto the molecular surfaces with an electronic density of 0.001 au. Gray, white, blue, red, and orange represent C, H, N, O, and P atoms, respectively.

Fig. S11

#### Table S1 | Specific activity of sample for XRF

| | Specific activity (U/mg) |
|---|---|
| $Zn^{2+}$-ShPPase | 66 ± 13 |
| $Zn^{2+}$-ShPPase for XRF | 31 ± 6 |

The metal content of $Zn^{2+}$-ShPPase was calculated based on the calibration curve (y=185.5x+59.418) with Zn standard solution (Fig. S8). The results indicated that the amount of zinc bound to the enzyme in the $Zn^{2+}$-ShPPase sample solution was 1.3 per molecule of enzyme. In addition, only the X-ray fluorescence spectrum of zinc was detected in the sample, indicating that only zinc, and not other non-target metals such as iron ions, is present in the active site. Furthermore, the activity of the $Zn^{2+}$-ShPPase sample for XAFS measurements and the fully activated $Zn^{2+}$-ShPPase sample were measured and compared. The fully activated $Zn^{2+}$-ShPPase sample was diluted in Activation buffer to be 0.5 mg/mL $Zn^{2+}$-ShPPase sample for XAFS measurement and left on ice for 2 hours to fully activate. As a result, the specific activity of the sample for XAFS measurement was 47% of that of the complete binuclear sample. Thus, 47% of the measured samples are binuclear, zinc ion-bound enzymes. Furthermore, since 1.3 zinc ions were bound per molecule, 36% were calculated to be uninuclear, and 17% to be without zinc ions. Therefore, it was determined that there were no free zinc ions in the sample solution that were not bound to the enzyme, which would be problematic for EXAFS analysis, due to the process of removing excess amounts of metal (ultrafiltration and spin columns).

#### Table S2 | Damage caused by X-ray irradiation

| | Specific activity (U/mg) |
|---|---|
| $Zn^{2+}$-ShPPase 1 scan | 45 ± 3 |
| $Zn^{2+}$-ShPPase 10 scan | 31 ± 4 |

Kinetic parameters determined from activity assays. Data are obtained with three times measurement.
